# Coupling between cerebrovascular oscillations and CSF flow fluctuation during wakefulness: An fMRI study

**DOI:** 10.1101/2021.03.29.437406

**Authors:** Ho-Ching (Shawn) Yang, Ben Inglis, Thomas M. Talavage, Vidhya Vijayakrishnan Nair, Jinxia (Fiona) Yao, Bradley Fitzgerald, Amy J. Schwichtenberg, Yunjie Tong

## Abstract

Cerebrospinal fluid (CSF) plays an important role in the clearance of metabolic waste products from the brain, yet the driving forces of CSF movement are not fully understood. It is commonly believed that CSF movement is facilitated by blood vessel wall movements (i.e., hemodynamic oscillations) in the brain. A coherent pattern of low frequency hemodynamic oscillations and CSF movement was recently found during non-rapid eye movement (NREM) sleep via functional MRI. However, questions remain regarding 1) the explanation of coupling between hemodynamic oscillations and CSF movement from fMRI signals; 2) the existence of the coupling during wakefulness; 3) the direction of CSF movement. In this resting state fMRI study, we proposed a mechanical model to explain the coupling between hemodynamics and CSF movement through the lens of fMRI. We found that the observed delays between hemodynamics and CSF movement match those predicted by the model. Moreover, by conducting separate fMRI scans of the brain and neck, we confirmed the low frequency CSF movement at the fourth ventricle is bidirectional. Our finding also demonstrates that CSF movement is facilitated by hemodynamic oscillations mainly in the low frequency range, even when the individual is awake.

## Introduction

The glymphatic system plays an important role in the clearance of metabolic waste products in the extracellular, interstitial space^1, 2^ and the distribution of homeostasis-sustaining compounds (e.g., glucose) in the brain^3^. Recent advancements in understanding the glymphatic system highlight its role in the development of neurodegenerative diseases^4–6^. The ventricles of the brain are a connecting network of cavities filled with cerebrospinal fluid (CSF). The ventricular system is composed of 2 lateral ventricles, the third ventricle, the cerebral aqueduct, and the fourth ventricle. The choroid plexuses of the ventricles produce CSF, which fills the ventricles and subarachnoid space. The circulation of CSF in the brain follows a cycle of constant production and reabsorption^7^. Unlike blood flow, which is driven by the heart pumping, CSF does not appear to have a single engine to generate flow^8^. CSF movement has been assumed to be mostly facilitated by the mechanisms associated with vessel wall movement, including vasomotion^9, 10^, pulsation^11^, inspiratory thoracic pressure changes^12, 13^, and/or osmotic pressure^14^.

Several magnetic resonance imaging (MRI) techniques have been used to investigate CSF flow in the brain, including 1) phase-contrast imaging^15^; 2) velocity density imaging^16^; 3) time-spatial labeling inversion pulse imaging^17^; and 4) 4D flow MRI^18^. These imaging techniques were often used to detect the direction of CSF motion within a single cardiac/respiration cycle. However, only a few studies reveal real-time CSF movement over a longer period or bulk flow in a wakeful state^11^.

Recently, Fultz et al. used a fast echo-planar imaging (EPI) sequence (TR<400ms) to study CSF flow^19^. They ingeniously placed the edge of the imaging volume (the first slice) at the fourth ventricle, which allows them to measure the flow of CSF using the inflow effect in the fMRI signal (rather than blood-oxygen-level-dependent (BOLD) contrast). They discovered a coherent pattern of oscillating electrophysiological, hemodynamic, and CSF dynamics that appears during non–rapid eye movement (NREM) sleep. Their results demonstrated the couplings between CSF movement and the hemodynamic signal, and low frequency electroencephalogram (EEG) during sleep. While still a qualitative approach to the measurement of CSF flux, the advantages of the fast EPI approach are obvious. First, the fast EPI sequence is widely available, and the data processing is straightforward. Second, through simultaneous recording, coupling between brain hemodynamics and CSF signal, which may be crucial in understanding the fundamental driving mechanism of CSF movement, can be examined. In this case,hemodynamics and CSF signal can be assessed by BOLD and in-flow contrast of fMRI, respectively.

While effective and innovate, the study by Fultz et al. is incomplete. First, they reported correlations between the derivative of the averaged fMRI signal in gray matter with CSF inflow signal (into the brain), assuming CSF dynamics in the fourth ventricle are driven by changes in cerebral blood volume (CBV). However, due to the limitation of the scan parameters, their method could only assess the inflow of CSF to the brain and not the outflow from the fourth ventricle (i.e., to the neck), We wondered if the same CBV-based mechanism could be demonstrated for CSF outflow. Second, the existence of CBV-CSF coupling was not fully explored during the awake state, when CBV changes may be driven primarily by non-stationary blood gases rather than with widespread electrical activity^20^.

This study was designed to address these questions. First, we proposed a simple mechanical model to explain the relationships between CSF movement and hemodynamic fluctuations in the fMRI signal. Second, we investigated the coupling between brain hemodynamics and CSF movement during the awake state by an fMRI scan. Last, an additional fMRI scan was employed to assess CSF outflow separately, and compare it with the brain hemodynamics. In the rest of the manuscript, we use “CSF movement” to represent the oscillations in CSF through the fourth ventricle, distinct from “CSF flux” which represents the bulk flow. We use the terms inflow and outflow to indicate the relative CSF movement towards brain and neck, respectively, with periodic events such as the cardiac and respiratory cycles, while remaining agnostic on the directionality of the CSF flux.

## Material and Methods

### Model

Even though the brain is not rigid, the skull forms a rigid “container” for the brain. As a result, the volumes of the constituents (blood, CSF, and brain parenchyma) consistently fluctuate to maintain the total volume needed for proper internal pressure—i.e., the Monro-Kellie doctrine^21^. According to this doctrine, the transfer of movements from the arterial walls into the surrounding tissue will eventually cause CSF to flow into and out of the spinal canal^22–24^. Multiple mechanisms cause arterial wall oscillations at various frequencies, such as cardiac pulsation (~1Hz) and vasomotion (<0.1Hz). They have all been shown to produce CSF movements^9–13^. However, the interpretation of the coupling between hemodynamic and CSF movement using fMRI signal is still not clear.

Here, we offer a general model that explains the coupling between fMRI hemodynamic signals and CSF movement. The model is based on the hypothesis that the cumulative effects of vessel dilations and contractions (pulsation, vasomotion) will exert force on the walls of ventricles (in which there are no blood vessels), especially the lateral ventricles, forcing CSF in and out of the fourth ventricle at the bottom (see Figure 1). In this model, the global mean of fMRI signals (GMS) is used as a surrogate signal to indirectly assess the cumulative effects of vessel dilations and contractions in the brain^20, 25, 26^. Notably, several previous studies had used this method for blood tracking^27–30^. In Figure 1, we illustrate volume changes in blood vessels and their consequential CSF movement observed at the fourth ventricle. In detail, the blood-rich region will expand because of vessel dilation. This will compress the volume of the lateral ventricle, forcing CSF out of the brain. The reverse happens when the blood vessels contract. However, the key point here is that CSF movement only happens at the transitions, during which the cerebral blood volume (CBV) changes (as shown by the arrows in Figure 1)^19^. When CBV is stable, no force is exerted on the ventricular wall, thus no flow of CSF. Therefore, the model predicts that CSF moves towards the neck through the fourth ventricle during CBV transition from low to high, and CSF move towards the brain during CBV transition from high to low.

**Figure 1.**
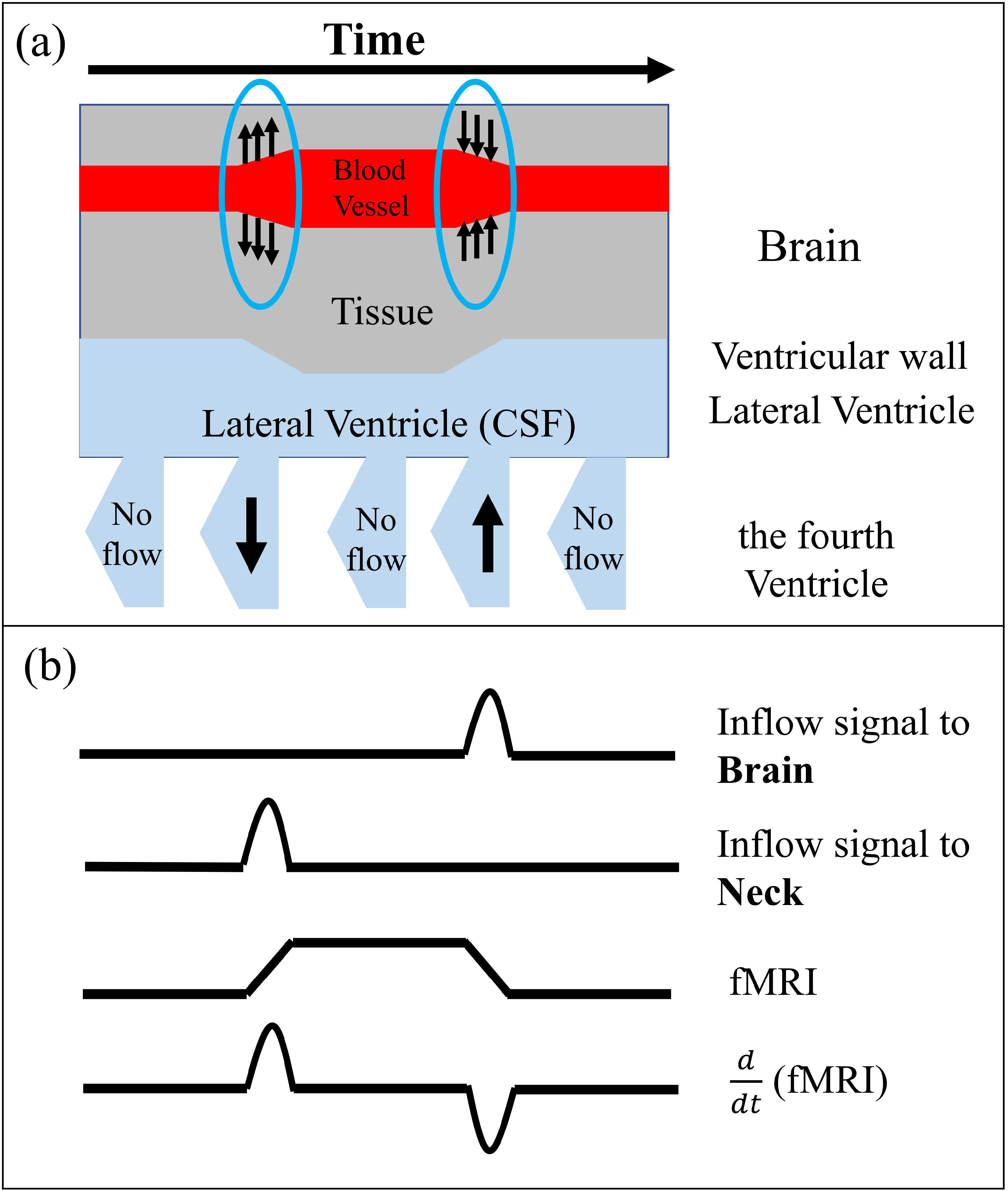
Model describes the derivative of fMRI signals are coupled with CSF Inflow and outflow fluctuations in the fourth ventricle. (a) Main structural of the model with relative brain (blood vessel and tissue) and ventricle structural (lateral and the fourth ventricle). Arrows circled with blue in brain show the vasodilation (arrows going out from blood vessel) and vasoconstriction (arrows going into blood vessel). Arrows in the fourth ventricle indicate the inflow and outflow fluctuation. (b) CSF inflow and outflow signal in the fourth ventricle affected by 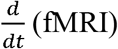 in the brain and time series in the brain 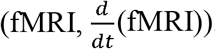.

Based on this model, we have several predictions (Figure 1): 1) CSF inflow (toward the brain) fluctuation should be negatively correlated with 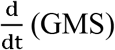 because the negative part of 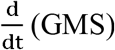 represents the contraction of CBV^19^; 2) CSF outflow (toward the neck) fluctuation should be positively correlated with 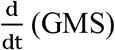 because the positive part of 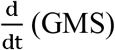 represents the dilation of CBV; 3) Since CBV changes are produced in compliance vessels, CSF flow fluctuation (in either direction) should lag behind the CBV change (i.e., 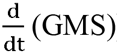) whether it is produced by neurovascular coupling or a blood-borne vasodilator such as CO_2_. To assess these predictions, we have conducted the following studies using specially designed scan locations and sequences.

### Experimental Design

#### Structural Scans

The study was approved by Purdue University’s Human Research Protection Plan (IRB-2019-303) and was conducted in accordance with application of Belmont Report principles (Respect for Persons, Beneficence, and Justice) and federal regulations at 45 CFR 46 and 21 CFR 50 and 56. Informed consent was obtained from all participants. Ten healthy individuals (5 female, 5 male, age range 23-30 years) were recruited. MRI data were obtained using a 3T SIEMENS MRI scanner (Magnetom Prisma, Siemens Medical Solutions, Erlangen, Germany) and a 64-channel head coil. Each participant underwent one T1-weighted, one T2-weighted, and two functional scans. For T1-weighted scans, Magnetization Prepared Rapid Acquisition Gradient Echo (MPRAGE) structural images were acquired with the following parameters (TR/TE: 2300/2.26 ms, 192 slices per slab, 321s, flip angle: 8 deg, resolution: 1.0mm X 1.0mm X 1.0mm). Parameters for T2-weighted scans were (TR/TE: 2800/409 ms, 208 slices per slab, 290s, resolution: 0.8mm x 0.8mm X 0.7mm). Both T1-w and T2-w images covered the head and the neck (Figure 2 (a)), to encompass the volumes of interest of the two fast EPI MRI scans (red and blue regions in Figure 2 (a, e)).

**Figure 2.**
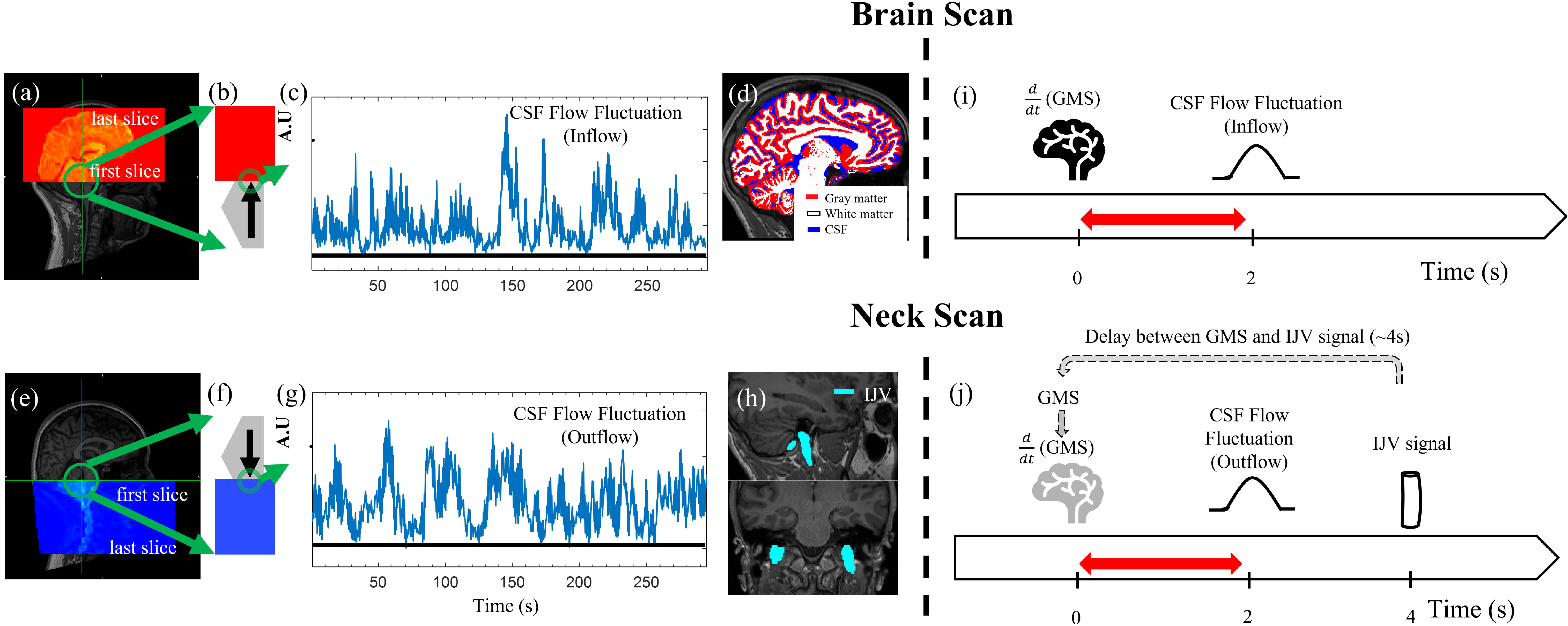
Low frequency oscillations in CSF signals in the fourth ventricle during wakefulness. Example scan positioning for (a) CSF brain scan and (e) CSF neck scan. The red shaded area in (a) and blue shaded area in (e) are the functional image coverages relative to the anatomy. The green circles point out the fourth ventricle. the fourth ventricle was enlarged as gray pentagons in (b) and (f) with arrows indicated CSF flow fluctuation direction and displayed with functional image (red and blue areas). Time series of a single CSF voxel from (c) brain scan and (g) neck scan. Both show large slow fluctuations at rest. (d) Brain segmentations (Gray matter, White matter, and CSF). (h) Blood vessels segmentations (IJV). (i) In the brain scan, CSF flow fluctuation (inflow to the brain) occurs ~2 s after 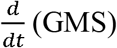. (j) In the neck scan, as there was no GMS measurement, IJV signal is used as a surrogate of GMS.

#### EPI scans

The functional resting state (RS) scans were acquired using a multiband echo-planar imaging sequence (FOV = 230 mm, acquisition matrix = 92 × 92, 48 slices, voxel size = 2.5 × 2.5 × 2.5 mm^3^, TR/TE = 440/30.6 ms, echo-spacing=0.51 ms, flip angle = 35°, hyperband acceleration factor = 8, multi-slice mode: interleaved). Participants were instructed to stay awake during the RS scans. CSF inflow fluctuation captured by a fast EPI sequence has been discussed in detail in Fultz et al.^19^. In short, as fresh fluid (that has not experienced radiofrequency pulses) move into the imaging volume, it induces higher signal intensity via apparent T1 shortening (i.e., inflow effect). To capture the inflow effect in both directions in the fourth ventricle, two 5-minute RS scans with different regions of interest were included. Both scans are positioned with the edge of the volume placed at the bottom of the fourth ventricle (see Figure 2 (a, e)), with one scan extending upward toward the top of the head (i.e., “brain scan”) and the other scan extending downward into the neck (i.e., “neck scan”). As a result, CSF inflow to the brain can be assessed by the brain scan, while CSF outflow from the brain can be assessed by the neck scan under the assumption that flow sensitivity in the opposite directions can be ignored due to steady-state excitation in those slices either above or below the slice of interest. The slice of interest, at the fourth ventricle, was consistently acquired first in the TR. The fourth ventricle was chosen for the following reasons: 1) From a physiological standpoint, it represents the conduit for CSF movement between the midbrain and neck. 2) Unlike the lateral ventricles, the fourth ventricle has a narrow shape which restricts the direction of the CSF flow (only allowing vertical movements). Therefore, the first EPI slice can be placed perpendicular to the flow direction to maximize the inflow effect. 3) We chose the middle part of the fourth ventricle over the other narrower parts to increase the likelihood of having some voxels (2.5mm) that are fully immersed in the CSF, with low partial volume from surrounding tissues.

### Data processing

#### CSF signals

All MR data were processed using FSL (FMRIB Expert Analysis Tool, v6.01; Oxford University, UK) and MATLAB. For analysis of CSF flow dynamics, a voxel in the fourth ventricle in the edge slice of the EPI data was identified with the help of the T1-weighted image (registered onto the EPI data). The time series of the voxel was extracted to represent CSF flow fluctuation in a certain direction. For CSF inflow fluctuation, the signal was extracted from the bottom EPI slice of the head scan as shown in Figure 2 (a, b). For CSF outflow fluctuation, the signal was extracted from the top slice of the neck scan as shown in Figure 2 (e, f). We hereafter refer to these two signals as “CSF inflow fluctuation” and “CSF outflow fluctuation”, even though they both stem from the inflow effect in fMRI.

#### Preprocessing

Since motion correction cannot be performed accurately on edge slices (given that tissue moves in and out of the imaging volume), only the slice-timing correction was performed prior to CSF signal extraction^19^. To make sure our observations were not due to motion artefacts, validations were performed which can be found in the supplemental materials (supplemental Table S5-7). Then, RS-fMRI data were preprocessed with the steps recommended by Power et al.: 1) slice-timing correction (FSL slicetimer) 2) motion correction (FSL mcflirt) and 3) spatial smoothing with a full width at half maximum (FWHM) of 5 mm isotropic Gaussian kernel was applied to the brain scan^31^. Spatial smoothing was not performed on the neck scan to avoid smoothing effect around the veins in the neck.

#### Data and statistical analysis

Segmentation was performed on the structural image. Several masks were made to investigate the coupling between CSF inflow fluctuation and fMRI signal from these segmentations (Figure 2 (d)). First, anatomical masks for gray matter, white matter, and CSF were made using an automated segmentation program (FSL fast)^32^. Second, we identified large veins in the neck (i.e., internal jugular veins (IJVs)) and made corresponding masks (Figure 2 (h)). In a previous RS-fMRI study, we found consistent positive correlation between the GMS of the brain and the fMRI signal from the IJVs (IJV signal lags the GMS by 4 seconds)^33^. Therefore, here we used the fMRI signal from the IJVs as a surrogate signal of the GMS (i.e., a delayed version) in the neck scan (Figure 2 (j)) Signals in IJVs were averaged as one joint IJV signal to improve the signal-to-noise ratio. More information can be found in the discussion.

After extraction, the fMRI signals including the CSF signal (from the fourth ventricle), the GMS from gray matter, white matter, CSF-regions (Figure 2 (d)), and signals from big veins (IJV, Figure 2 (h)) - were first linearly detrended and then band-pass filtered (0.01 to 0.1 Hz) using a zero delay, fourth-order Butterworth filter to extract the low frequency oscillations (LFO). Cross correlations (MATLAB xcorr, lag range = ± 45 seconds) were calculated between (1) 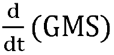 and CSF inflow fluctuations for the head scan, and (2) 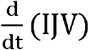 and CSF outflow fluctuations for the neck scan. The maximal cross correlation coefficients (MCCC) and the corresponding delays were calculated for all the research participants. One-sample Kolmogorov-Smirnov test was used to test the distribution of MCCCs. Since our results are not normally distributed, one-sample Wilcoxon signed rank test (one tail) against 0.3 (for testing positive MCCC) or −0.3 (for testing negative MCCC) was applied on MCCCs to minimize spurious correlation, which were calculated between (1) 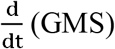 and CSF inflow fluctuations, and (2) 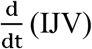(IJV) and CSF outflow fluctuations for all the research participants. This approach was applied based on a previous fMRI study which found significant (p<0.008) MCCC threshold of ±0.3 by applying randomized test with 1000 iterations within 100 healthy adults^34^. Notably, a significant (p<0.01) MCCC threshold of ±0.28 was also found in another study using a similar method^35^. Moreover, to assess the coupling between CSF inflow fluctuation with the whole brain and understand any spatial characteristics of this coupling, the CSF inflow fluctuation was cross-correlated with 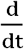 (fMRI signal) for every voxel in the brain (except the lateral ventricles). The MCCC and delay maps were derived for each participant.

Although we used a fast EPI scan with TR=440 ms, it is possible that the dynamics we observe are caused by aliasing of the cardiac cycle. To investigate pulsatile oscillations, we extracted the full cardiac pulsation waveform from the CSF signal and the brain fMRI scan from an example participant using Happy, which is a method to estimate cardiac pulsation from raw fMRI without additional physiological measurements^36^. The coupling between the CSF and the brain signals at cardiac frequency were assessed. Lastly, the CSF signal powers in cardiac and low frequencies were compared.

## Results

### CSF flow fluctuation during wakefulness

Figure 2 (c) shows an example of CSF flow fluctuation obtained from the brain scan. First, it shows that during an awake state, CSF inflow fluctuation is clearly detectable in agreement with Fultz et al. for deep sleep^19^. Second, similar fluctuation patterns were detected in the neck scan (Figure 2 (g)), which indicates the existence of a similar CSF outflow fluctuation. It is worth noting that the CSF flows are unidirectional, with flat baseline after detrending (black lines in Figure 2 (c, g)).

### CSF flow fluctuation and 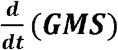

Figure 3 displays example low frequency data from one research participant. It shows 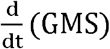 of the brain and CSF inflow fluctuation in (a-c), and the same participant’s 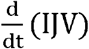 and CSF outflow fluctuation in (d-f). As shown in Figure 3, CSF inflow fluctuation matched the lower part of band-pass filtered 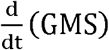 in (b) (CSF inflow fluctuation was flipped for demonstration; MCCC= −0.75), while CSF outflow fluctuation matched the upper part of band-pass filtered 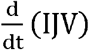 in (e) (MCCC=0.84). Analyzing cross-correlation coefficients at different delays reveals that from the delays, CSF inflow fluctuation lagged 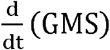 by about 2.2s (Figure 3 (c)) and CSF outflow fluctuation led 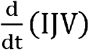 by about 2.2s (Figure 3 (f)).

**Figure 3.**
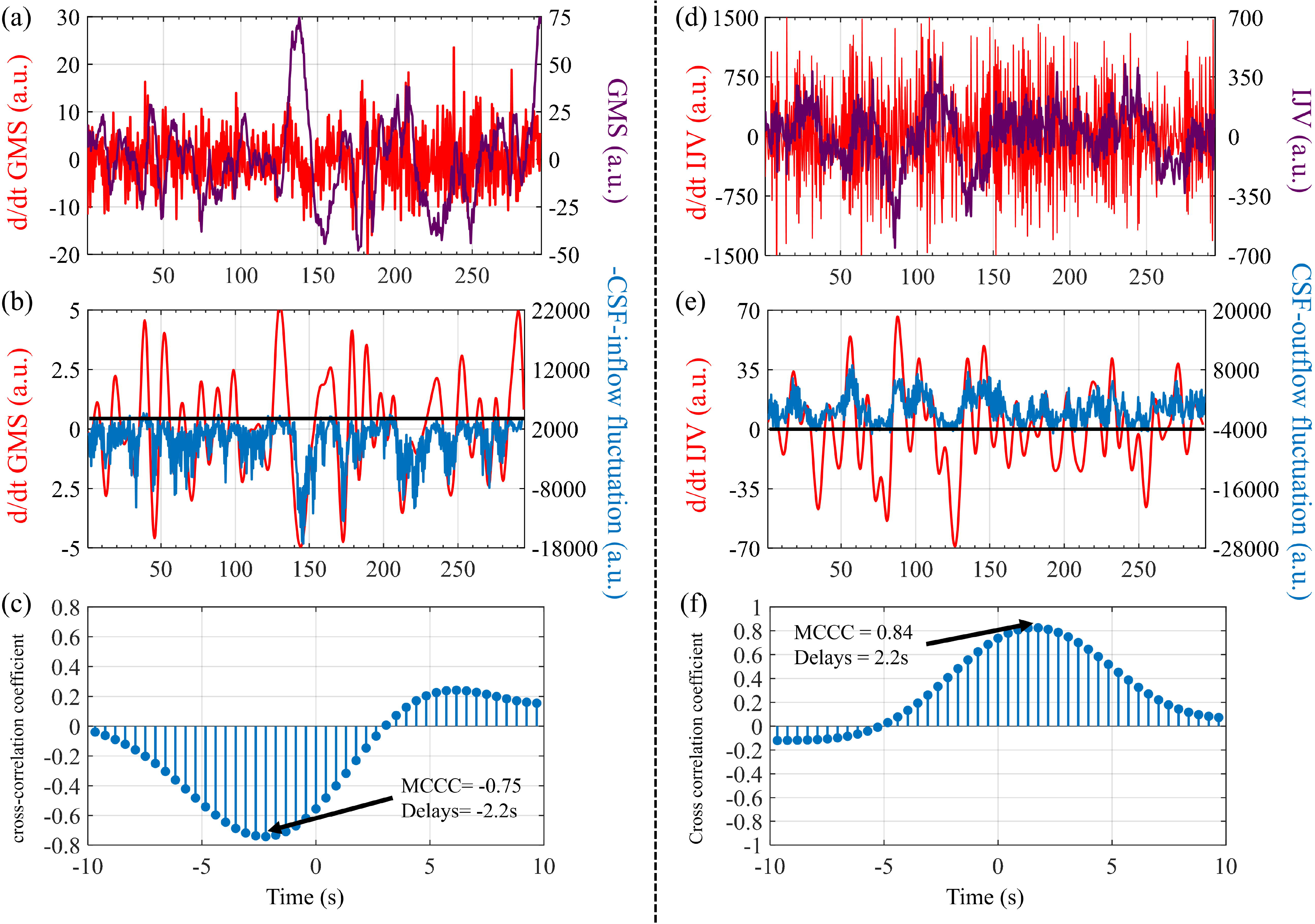
The relationships between global mean signal (GMS) of the brain and CSF inflow fluctuation, and IJV signal (neck) and CSF outflow fluctuation from one research participant. (a) Global mean signal (GMS) and derivative of global mean signal 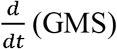). (b) Band-pass filtered 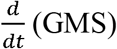 and negative CSF inflow fluctuation. (c) Cross-correlation between 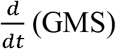 and CSF inflow fluctuation. (d) IJV signal (IJV) and derivative of IJV signal 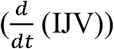. (e) Band-pass filtered 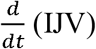 and CSF outflow fluctuation. (f) Crosscorrelation between 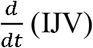 and CSF outflow fluctuation.

Figure 4 (a) shows the individual and average results of MCCCs and delays calculated between 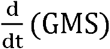 and CSF inflow fluctuation from the brain scans. Also, Table 1 presents all cross-correlation results (cross-correlation coefficients and delays). Here 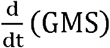 was calculated separately for the gray matter, white matter, and CSF regions in the brain. The MCCCs from all the participants are high for all the tissue types (MCCC for gray matter: −0.76±0.07 (p<10^−3^); for white matter: −0.69+0.10 (p<10^−3^); and for CSF: −0.76±0.07 (p<10^−3^)). The corresponding delays are around −2s—i.e., 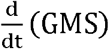 signal leads the CSF inflow fluctuation in fourth ventricle. Among the three tissue types, 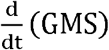 from gray matter, white matter, and CSF-region led CSF inflow fluctuation by 2.20±0.55s, 2.02 ± 0.69s and 1.36 ± 0.60s, respectively. Figure 4 (b) shows the average results of MCCCs and delays calculated between 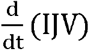 and CSF outflow fluctuation. The averaged MCCC is 0.73 ±0.11 (p<5 x 10^−3^) (participant 6 was excluded as an outlier) with average delay of 1.66+0.87s. As we mentioned previously, after adjustment for the delay between the fMRI signal of the IJV and the GMS (total ~4s), CSF outflow fluctuation should lag the brain signal 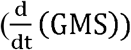 by about 2s^28^ (as illustrated in Figure 2 (i, j)).

**Figure 4.**
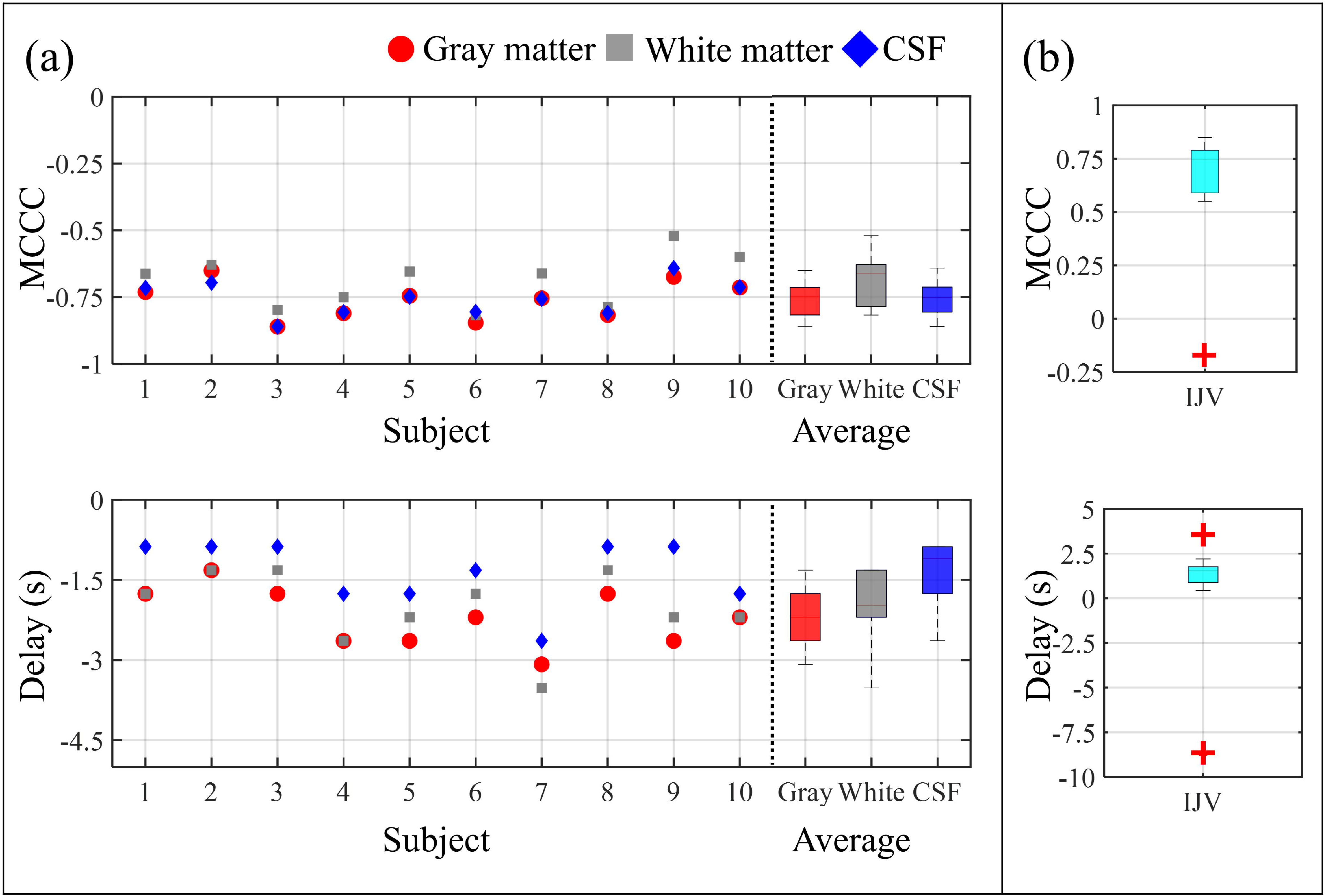
The summary of cross-correlations between CSF inflow fluctuation and 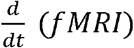 in different region of interests (ROIs). (a) MCCCs and delay times between CSF inflow fluctuation and 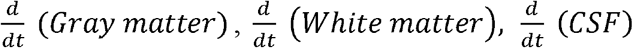 in each subject and in all subjects. All the measured 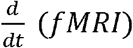 in different ROIs is ahead of CSF inflow fluctuation with high MCCCs. (b) MCCCs and delay times between CSF outflow fluctuation and 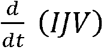. Most of the CSF outflow fluctuation is ahead of 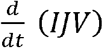 with high MCCCs.

**Table 1.** Cross-correlation results between 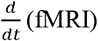 and CSF flow fluctuations, the negative lag means 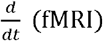 is leading CSF inflow signal. Red highlights the outliers that were excluded from the further calculation. *: cross-correlation between 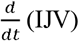 and CSF outflow signal was calculated without outlier (participant 6).

|  | Gray matter |  | White matter |  | CSF |  | IJV |  |
| --- | --- | --- | --- | --- | --- | --- | --- | --- |
| participants | xcorr | lag (s) | xcorr | lag (s) | xcorr | lag (s) | xcorr | lag (s) |
| 1 | -0.73 | -1.76 | -0.66 | -1.76 | -0.72 | -0.88 | 0.64 | 3.52 |
| 2 | -0.65 | -1.32 | -0.63 | -1.32 | -0.70 | -0.88 | 0.72 | 1.76 |
| 3 | -0.86 | -1.76 | -0.80 | -1.32 | -0.86 | -0.88 | 0.84 | 2.20 |
| 4 | -0.81 | -2.64 | -0.75 | -2.64 | -0.81 | -1.76 | 0.79 | 0.44 |
| 5 | -0.74 | -2.64 | -0.65 | -2.20 | -0.75 | -1.76 | 0.78 | 1.76 |
| 6 | -0.85 | -2.20 | -0.82 | -1.76 | -0.81 | -1.32 | -0.17 | -8.80 |
| 7 | -0.75 | -3.08 | -0.66 | -3.52 | -0.76 | -2.64 | 0.59 | 1.32 |
| 8 | -0.82 | -1.76 | -0.79 | -1.32 | -0.81 | -0.88 | 0.85 | 1.32 |
| 9 | -0.67 | -2.64 | -0.52 | -2.20 | -0.64 | -0.88 | 0.55 | 1.76 |
| 10 | -0.71 | -2.20 | -0.60 | -2.20 | -0.71 | -1.76 | 0.77 | 0.88 |
| mean | -0.76 | -2.20 | -0.69 | -2.02 | -0.76 | -1.36 | 0.73* | 1.66* |
| std | 0.07 | 0.55 | 0.10 | 0.69 | 0.07 | 0.60 | 0.11* | 0.87* |

### Spatial-temporal information of the coupling of CSF inflow with the brain signal

Figure 5 shows the averaged MCCCs and delay maps derived from the voxel-wise cross correlation between CSF inflow fluctuation and 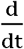 (fMRI signal). Negative MCCCs were found in most of the tissue-containing voxels with distributions shown in Figure 5 (a). Regions with the highest negative MCCCs are found in the gray matter, especially in high blood density regions such as the visual cortex. It is worth noting that the positive correlations (shown in cool colors) were found in the voxels next to the lateral ventricles (Figure 5 (a, b)), likely due to partial volume effects.

**Figure 5.**
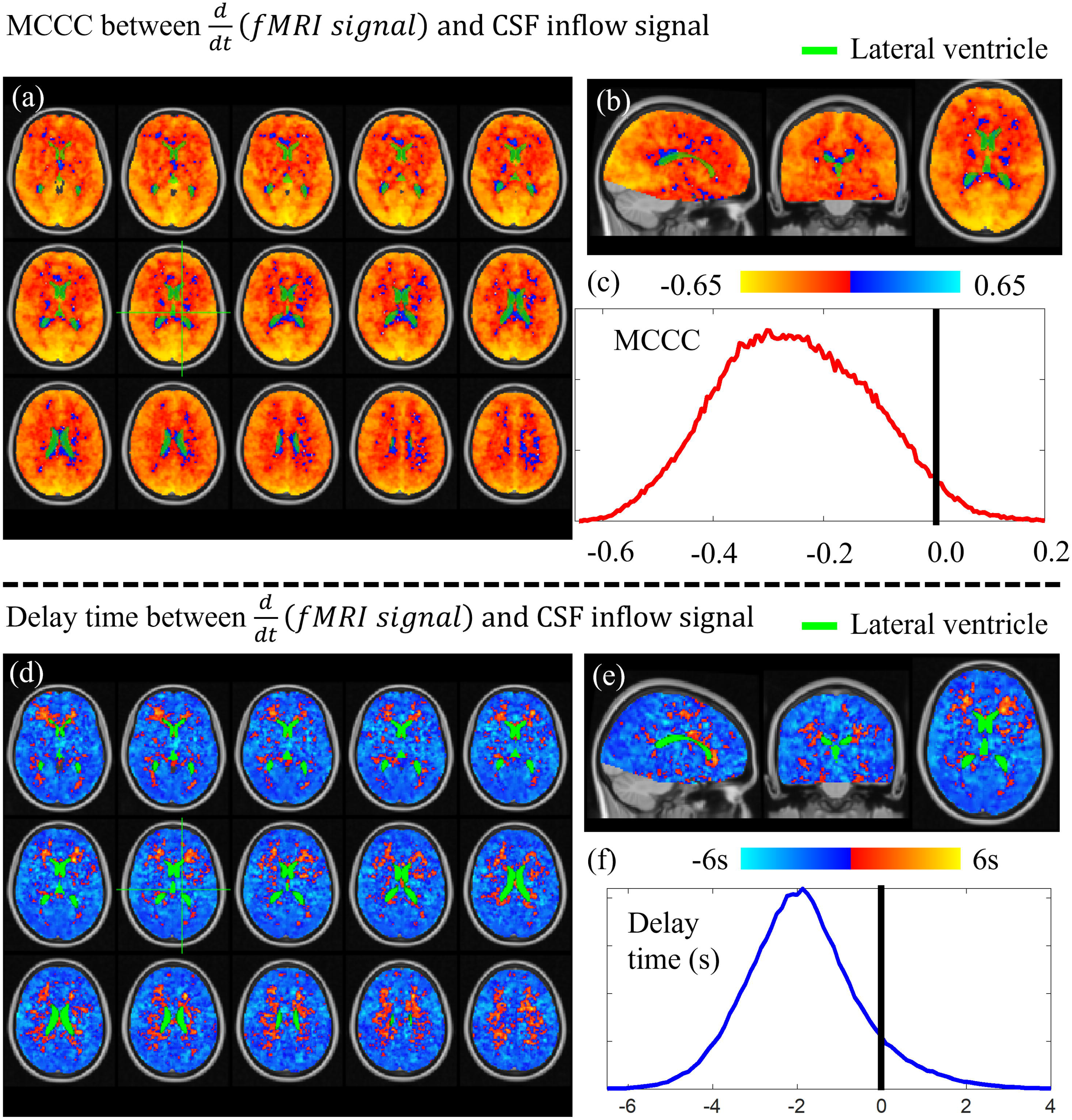
Group average of voxel wise cross-correlation between 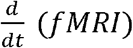 and CSF inflow fluctuation. (a) Lightbox view of whole brain MCCCs shows that most of the 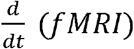 in the brain were negatively (red-yellow) correlated with CSF inflow fluctuation. (b) Whole brain MCCCs in sagittal, coronal and transverse views. (c) Distribution of MCCC between time series in whole brain voxels and CSF inflow fluctuation (d) Lightbox view of whole brain delay times shows that most of the 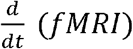 is ahead of CSF inflow fluctuation, (e) Whole brain delay times in sagittal, coronal and transverse views. (f) Distribution of delay time between time series in whole brain voxels and CSF inflow fluctuation. Green masks are used for the lateral ventricle.

Negative delays were found in most of the voxels (cold colors in Figure 5 (d, e)) with distributions shown in Figure 5 (f). Negative delays indicate that 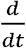(*fMRI signal*) is leading CSF inflow fluctuation. In contrast, few voxels contain positive delays (warm colors in Figure 5 (d, e)), mostly located in white matters. This result is consistent with the findings in Figures 3 and 4.

### Cardiac pulsation signal found in CSF fluctuation and the brain signal

Figure 6 (a) is example data of CSF inflow fluctuation taken from one participant’s head scan and its cardiac component extracted from fMRI data without additional physiological measurement^36^ (lower panel). To compare the cardiac components extracted from CSF fluctuation to that from the brain, small sections of both time series were displayed in Figure 6 (b). There is no clear time delay between cardiac components extracted from CSF fluctuation and that from the brain. To compare the effect of pulsation and slow oscillation on CSF flow fluctuation, power spectrum of the signal was shown in Figure 6 (c), in which the heartbeat signal (~0.87Hz) is much smaller than that of LFOs, suggesting negligible contribution of the cardiac cycle on our data.

**Figure 6.**
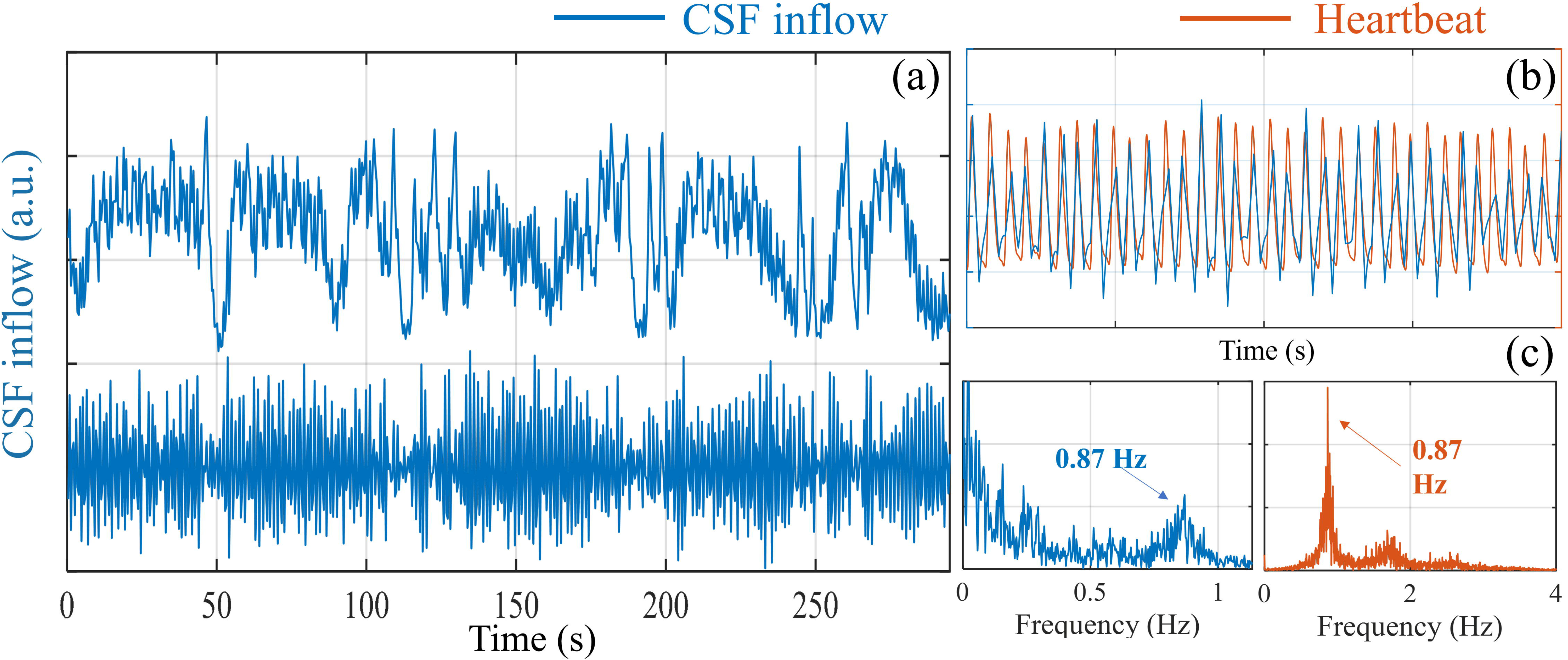
Cardiac pulsation signal found in CSF fluctuation. (a) CSF inflow fluctuation (upper panel: raw data of CSF inflow; lower panel: high-passed CSF inflow fluctuation (>0.6Hz)). (b) Segment of High passed CSF inflow fluctuation (>0.6Hz, blue) and heartbeat signal (orange) derived by Happy, which is a method to estimate cardiac pulsation^36^. (c) power spectrum of high frequency CSF inflow fluctuation (>0.6Hz, left) and power spectrum of heartbeat signal estimated by Happy (right)^36^.

## Discussion

This study used a simple model to explain the correlation and delays between 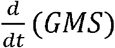 of fMRI and the CSF fluctuation signal measured at the fourth ventricle. With both brain and neck scans, we verified that CSF movement at low frequencies were bi-directional and found that coupling between the brain hemodynamics and CSF flow fluctuation exists for awake participants in the supine position.

### CSF flow fluctuation detected during wakefulness

Figure 2 shows that CSF inflow and outflow fluctuations were captured by the brain and the neck scans, respectively, during wakefulness. Since the CSF signal originates from an inflow effect that is sensitive to CSF flow in one direction (i.e., into the scan volume), a brain scan can only capture the CSF inflow fluctuations while a neck scan can only capture the CSF outflow fluctuations. The flat sections found in both Figure 2 (c, g)) indicate the time periods when the CSF flow was in the undetectable direction (e.g., CSF outflow during the brain scan). Two important things can be derived from this observation. First, low frequency CSF movement is bi-directional, like pulsatile CSF movement^2, 9, 37^ Therefore, our measurement only reflects the flow fluctuation, instead of CSF flux or bulk flow. Second, as stated by the model, the 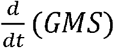 is sensitive to the transitions of CBV in the brain, and change of CBV is likely to be the driving force of the CSF movement. Unlike the CSF signal which can only report the flow fluctuation in one direction, the 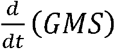 captures the full dynamic of the CBV changes. That is, the positive part of 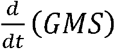 represents CBV dilation (leading to CSF outflow), while the negative part of 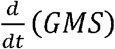 represents CBV contraction (leading to CSF inflow). These predictions were validated in this study as showed in Figure 3 (b, e), where we found the positive and the negative parts of 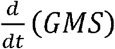 matched well with CSF outflow and inflow fluctuations, respectively. This study explains the internal relationship between 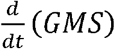 and CSF movement in both directions at the fourth ventricle.

### CBVand GMS (IJV signals)

It has been shown that during resting state, ongoing dynamics in the vasculature which are independent of neural activity are likely due to vessel-autonomous or respiratory-related oscillations in the arterial diameter(~30%)^31, 38, 39^. CBV_a_ (arterial CBV) fluctuations have been found in arteries throughout the brain^40–43^. In our previous research, these oscillations were even found in the internal carotid arteries (ICA) earlier than their appearances in the brain^33, 34^. These upstream oscillations move along the arterial tree into the brain and cause corresponding CBF changes in the downstream capillaries, venules, and veins, following the path of blood flow. Animal studies have confirmed that these vasomotor oscillations in the arteriole diameter drive oscillations in the rate of brain tissue oxygenation^44, 45^, which then leads to delayed passive dilations in the draining venules^46, 47^. These CBF related oxygenation changes in capillaries, venules, and veins, induce the BOLD contrast that we observed as the GMS. The same CBF changes also lead to passive dilations in the draining venules and veins^48^ with smaller scale in vessel diameters^49, 50^. Therefore, the GMS reflects the low frequency CBV changes in both arterioles and venules, primarily by modulation of CBF that results in the well-known dilution model of BOLD^20^.

The same CBF changes will later be observed via BOLD signal in large draining veins, such as the superior sagittal sinus (SSS) and IJV, as the blood moving out of the brain. For example, a previous study with 90 resting-state scans showed the fMRI signal in IJV lagged that of SSS by 1s, while the fMRI signal in SSS lagged the GMS by 3s^33^. This is consistent with a blood flow phenomenon mediated upstream by changes in CBV. For further validation, we rescanned two participants using a similar protocol but with slices covering from the brain to the neck (TR increased to 1s). We confirmed that the LFO in the IJV is correlated with the GMS (participant 1: 0.46; participant 2: 0.69) with few second delay (participant 1: 3.39s; participant 2: 4.52s). The results and scan protocols can be found in the supplemental material (Supplemental Figure S3). Based on these findings, we also confirmed that the IJV signal can be used as a delayed surrogate signal of the GMS in the neck scan, as illustrated in Figure 2(j), in which the actual GMS could not be obtained.

### Time delays

Based on the results of the brain scan (Figure 4(a)), 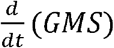 was found to lead the CSF inflow fluctuation measured at the fourth ventricle by about 2s. A delay was predicted by the model, indicating CBV changes are a candidate mechanism for CSF flow fluctuation, though maybe not the only one, as we discuss in a separate section below. Similar results in delay were also shown in Figure 4(b). However, the CSF outflow fluctuation leads 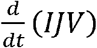 signal by about 1.62 s. From previous studies, the fMRI signal of IJV lags the GMS by about 4 seconds^33^. After adjustment for this delay, CSF outflow fluctuation should still be lagging the true 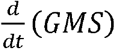—i.e., the brain signal—by about 2 seconds, which is consistent with the findings in the brain scan (illustration in Figure 2 (j)).

The time delays differ from the results obtained from the cardiac pulsation (Figure 6(b)). As cardiac pulsation travels through the brain (i.e., fast pressure wave), it affects all the brain regions almost simultaneously. This synchronized oscillation will place pressure on the ventricular wall and leads to instantaneous CSF flow fluctuation at the fourth ventricle, as first predicted by the Monro-Kellie doctrine^21^. Adams et al. has measured the intracranial volume dynamics using Displacement Encoding with Stimulated Echoes (DENSE)^51^. They demonstrated the dynamic balance between CSF flow and brain tissue volume over one cardiac cycle. Interestingly, they also found a small delay between peak gray and white matter volumetric strain (within one cardia cycle), which, they attributed to the perfusion from gray matter to white matter following the cerebrovasculature. From our study, we believe CBV also oscillates at a much lower frequency (0.01-0.1Hz), which leads to the low frequency CSF flow fluctuations. Importantly, the low frequency CBV changes do not move as fast as cardiac pulsation. The pressure induced by CBV propagates through the vasculature as blood flow, which takes seconds. The spatial-temporal pattern of the low frequency wave moving through the brain has been validated in several groups of human subjects^30, 52^. Similarly, various delays (0.5-6s) were also found between the CBV signal primarily from arterioles and the BOLD signal from individual venules^48, 53^ in animal studies.

As we consider the changes in CBV 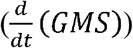 as a cause for “pressure waves”, the corresponding delay map (Figure 5(d)) represents the propagation of these waves in the brain. As this low frequency pressure wave moves through the brain (warm colors in Figure 5(d)), it may influence the CSF flow only when it reaches the vicinity of lateral ventricles or vascular territories, which is relatively late in the propagation path. Like the cardiac pulsation propagation, it moves from gray matter to white matter (see Figures 4 and 5). As mentioned above, the GMS represents the BOLD effect from capillaries, venules, and veins in the brain, but mostly from capillaries. The CBF change that produces the dilution of deoxyhemoglobin on the venous side implies a concomitant CBV change. These delayed passive dilations in the venules and veins that surrounds the lateral ventricles, might directly cause the CSF movement at the fourth ventricle. This would explain the delays (~2s) between 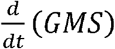 and CSF flow fluctuations. However, a few observations in Figure 5 do not fit the proposed model. First, the voxels with the positive delays (warm colors in Figure 5 (d)), which cover much of the extended region from the ventricles into the white matter, do not overlap with the positively correlated voxels in Figure 5 (a). Second, the delay values are positive, indicating they lag a few seconds after CSF inflow fluctuation. We do not fully understand the reason for this. It might indicate there are secondary interactions between CSF inflow and white matter, such as an elastic rebound or some other motion-related phenomenon in the tissue.

### Various physiological mechanisms for CSF flow or flow fluctuation

Previous studies have proposed several driving forces for CSF flow, which included 1) cardiac pulsation^54^, 2) vasomotion^10^, and 3) respiration^13, 37^. Some studies demonstrate that pulsation is a driving force for CSF^2, 54, 55^. Others studies used mathematical model to argue that arteriolar pulsation alone are too weak to drive CSF circulation or flux^56, 57^. However, the pulsation might still contribute to the fast exchange of CSF-interstitial fluid (ISF) through mixing and diffusion^51^. This could be the crucial first step in removal of the waste products in the brain.

Figure 6 shows that, compared to cardiac pulsation, LFOs are dominant in the CSF signal. However, in this study, we can only measure CSF flow fluctuations in one direction at a time in each scan and cannot determine the CSF flux directly. We estimated the CSF flux combining two consecutive scans (brain and neck scans) and found no clear flux during the short scan time (see Supplemental Figure S1). Many animal studies have tried to understand the CSF flux using contrast agents. However, the pathways for the lymphatic drainage of CSF are still not clear^58^.

We showed that low frequency CSF movement was detected in both directions. Like the effect of cardiac pulsation^9, 51^, the low frequency CSF flow fluctuations might also contribute to the fast exchange of CSF-ISF through mixing and diffusion. It would be of great interests to understand the distinct roles played by bi-directional CSF flow fluctuation and CSF flux in the clearance of the wastes in the brain^1, 8, 59^. Nevertheless, increased low frequency vasomotion (0.1 Hz) in arterioles has been found to increase clearance rate from perivascular space^10^.

In addition to pulsation and LFOs, several lines of work have shown that respiration is instrumental in driving CSF through the aqueduct by using real-time phase contrast MRI measurements^11–13, 37, 60^. Furthermore, Dreha-Kulaczewski et al. proposed that increased CSF flow from caudal to cranial during inspiration was considered compensation of venous blood leaving from head due to lower intrathoracic pressure^13^. Based on this theory (i.e., pressure), the CSF response to respiration should be almost instantaneous. However, no respiration belt was used in this study, which limited us from assessing the respiratory effect on the CSF flux or flow fluctuation (see limitations of the study).

Our study focused on the LFO of the signal, not the respiration frequency. The natural breathing frequencies of our participant were from 0.2-0.4 Hz (The power spectra of the CSF inflow/outflow and the GMS signals of all the participants can be found in the Supplemental Table S1, S2). Like cardiac pulsation, the power of the respiration signal is much less than that of the LFOs, indicating that the arterial LFO is the dominant force in CSF flow fluctuation, at least in resting state without using any forced/paced breathing protocol. In addition, forced or paced breathing as used in Dreha-Kulaczewski’s study might invoke different physiological responses/mechanisms, other than the ones under regular breathing^13^.

### Sleep vs. awake states

Previous research documents increases in the cortical interstitial space by more than 60% during sleep when compared with an awake condition, resulting in efficient convective clearance of *β*-amyloid and other compounds^1^. This highly sleep-dependent clearance was observed in both human and mouse models^61, 62^. In our study, we demonstrated a strong coupling between brain hemodynamics and CSF flow fluctuation in awake participants. It suggests that low frequency fluctuations in CSF motion are maintained during the waking hours. The fMRI-signal-to-CSF delays we measured in awake subjects (~2 sec) are very close to the delays found by Fultz et al. for subjects in deep sleep (1.8 sec). Despite similar delays, however, the periods between CSF fluctuations detected in deep sleep by Fultz et al. are much longer than the breathing rate and suggest a neural origin. In our prior work, we found that LFOs in awake subjects can be explained by changes in respiration rate and depth, suggesting an origin in blood gases in the awake case; most likely the pCO_2_. Thus, the transition from awake to deep sleep may involve a shift in the principal cause of CBV oscillations in the brain. It would be of great interest to understand the fundamental differences between sleep and the wakeful states, in term of vasomotion, CSF flux, flow fluctuation, coupling, and overall efficiency of clearance of the inflammatory proteins and metabolites.

### Limitation and future studies

There are several limitations to the study which we hope to address in future experiments. First, physiological parameters (e.g., respiratory, cardiac and P_ET_CO_2_ measurements) have not been recorded, which limited us from assessing their relationships with LFOs in CSF and BOLD fluctuations. Second, we did not fully sample the cardiac signal in the fMRI data due to the low temporal resolution, thus we cannot systemically assess the relationship between the GMS and the CSF signals associated with cardiac pulsation. Third, we did not collect simultaneous EEG data. Thus, we could not confirm if the neuro-vascular coupling (demonstrated in the Fultz et al^19^) works in the wakeful state. In future studies, we will incorporate respiration (chest belt, P_ET_CO_2_), cardiac pulsation and EEG measurements and increase the temporal resolution of fMRI. Moreover, we will compare regular and forced breathing, to understand the differences in the underlying mechanisms.

## Conclusions

A model, based on the Monro-Kellie doctrine^21^, was proposed in this study to interpret fMRI signal and link fMRI signal with certain brain dynamics. Two fMRI scans were conducted to validate the model and several predictions. We found 1) coupling existed between LFOs of 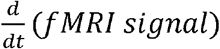 and CSF flow fluctuations when participants were awake; 2) the LFOs of 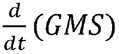 occurred about 2.2 seconds earlier than those of CSF fluctuation at the fourth ventricle; 3) low frequency CSF movement at the fourth ventricle is bi-directional. Together, we conclude that the arterial LFO is the dominant force in driving CSF flow fluctuation during wakefulness. These findings can help understand the mechanics of CSF flow fluctuation and develop new interventions to increase the clearance rate in the brain, especially for patients with neurodegenerative diseases.

## Acknowledgements

We would like to thank Drs. Blaise B. Frederick, Ahmed A. Khalil, Lia M. Hocke, Ioannis Pappas, and Xiaopeng Zhou for useful discussion. We would also like to thank Ms. Antonia Susnjar for data collection. Lastly, we would like to thank two anonymous reviewers. The manuscript has been improved significantly with their comments. This work was supported by the National Institutes of Health, Grants S10 grant (S10 OD012336 - 3T MRI Scanner dedicated to Life Sciences Research PI: Ulrike Dydak). Indiana CTSI Core Facility Grant.

## Author contribution

Yunjie Tong, Ho-Ching (Shawn) Yang, and Ben Inglis conceived of the presented idea and developed the theory. Ho-Ching (Shawn), Vidhya Vijayakrishnan Nair and Jinxia (Fiona) Yao performed the computations and verified the analytical methods. All authors provided critical feedback and helped shape the research, analysis and manuscript.

