## Supplemental materials for "Coupling between cerebrovascular oscillations and CSF flow fluctuation during wakefulness: An fMRI study"

**Respiration**

As we mentioned in the main manuscript, several real-time phase contrast MRI studies indicated that respiration is crucial in moving CSF through the aqueduct ^1-5^. Respiration was investigated by accessing power spectrum for CSF flow fluctuation (brain/neck), Global mean signal and IJV signal (Table S1-2). For CSF inflow fluctuation (brain), only two out of ten participants (participant 3 and 6, Table S1) show relatively large respiratory signal (0.2-0.4 Hz). Notably, for CSF outflow fluctuation (neck), eight out of ten participants (participant 2-7, and 9-10, Table S1) show large respiratory signal. These results indicate CSF outflow fluctuation tends to be influenced by respiration. For GMS (brain) and IJV signal (neck), respiratory signal can only be found in part of the participants. For GMS, four out of ten participants (participant 6-7 and 9-10, Table S2) show relatively large respiratory signal (0.2-0.4 Hz). For IJV signal, four out of ten participants (participant 4, 6, 7, and 10, Table S2) show large respiratory signal. However, further examinations are necessary to understand the respiratory influence on fMRI signal.

**Derivative caused leading** $\frac{\mathbf{d}}{\mathbf{dt}}\mathbf{(GMS)}$ **to CSF inflow fluctuation?**

A scenario was considered in this study: the $\frac{d}{\mathrm{dt}}(GMS)$ leading the CSF inflow fluctuation could be caused by derivative operation. To disprove, cross-correlation was applied to calculate the time delay between $\mathrm{GMS}$ and CSF inflow fluctuation. However, inconsistent correlations were found ($\bar{\mathrm{MCCC}}: -0.08 \pm0.66; \bar{\mathrm{Delay}}: -2.64\pm5.24 (s)$). The $\mathrm{GMS}$ and CSF inflow fluctuations were either positively correlated with negative delay or negatively corelated with positive delay (Table S3).

Furthermore, cross-correlation was applied to calculate the time delay between $\mathrm{GMS}$ and the $\frac{d}{\mathrm{dt}}(GMS)$ to see if the derivative operation brings the resulting signal ($\frac{d}{\mathrm{dt}}(GMS)$) forward to the original signal ($\mathrm{GMS}$). Our results show the averaged MCCC and time delay between $\frac{d}{\mathrm{dt}}(GMS)$ and $\mathrm{GMS}$ were low with high standard deviation ($\bar{\mathrm{MCCC}}: -0.015 \pm0.71; \bar{\mathrm{Delay}}: -0.13\pm4.44 (s)$). The $\frac{d}{\mathrm{dt}}(GMS)$ and $\mathrm{GMS}$ were either positively correlated with negative delay or negatively corelated with positive delay (Table S4). Together, $\mathrm{GMS}$ is periodic for both $\frac{d}{\mathrm{dt}}(GMS)$ and CSF inflow fluctuation, which results in inconsistent MCCC and time delay. The derivative operation cannot bring $\frac{d}{\mathrm{dt}}(GMS)$ forward to CSF inflow fluctuation.

| participant | Spectrum for CSF inflow fluctuation (brain) | Spectrum for CSF outflow fluctuation (neck) |
| --- | --- | --- |
| 1 | 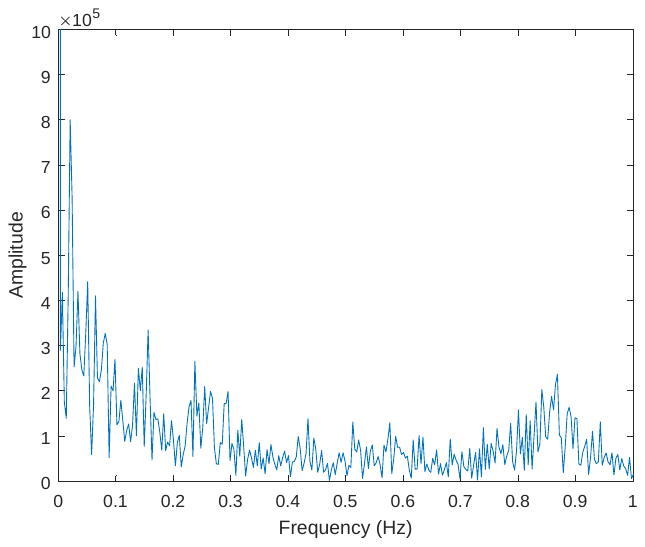 | 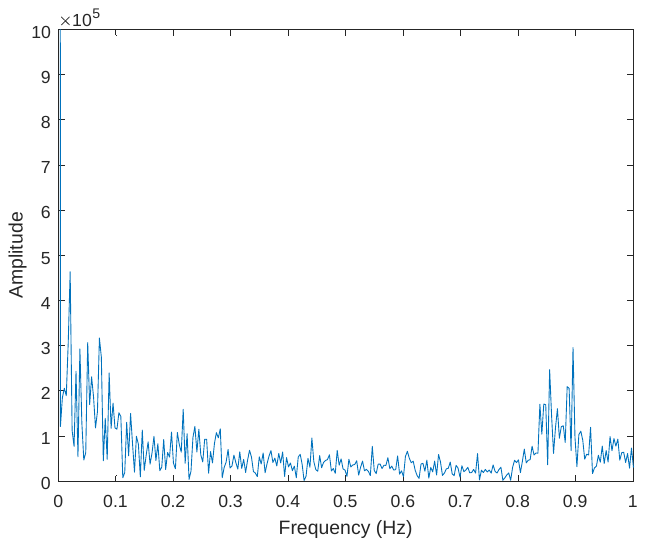 |
| 2 | 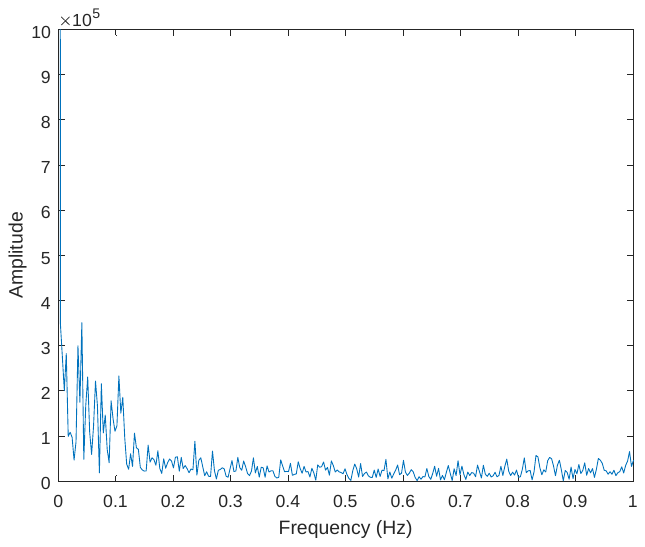 | 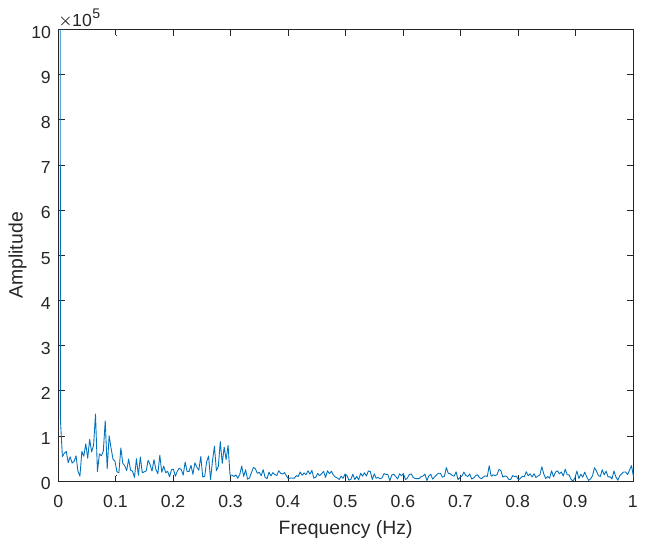 |
| 3 | 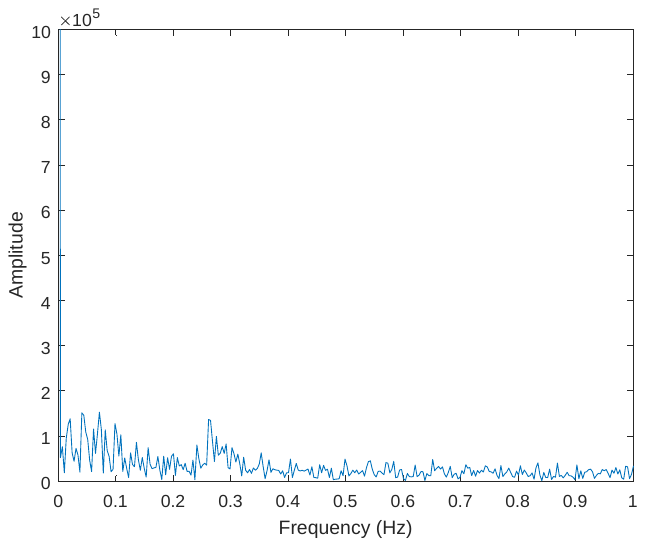 | 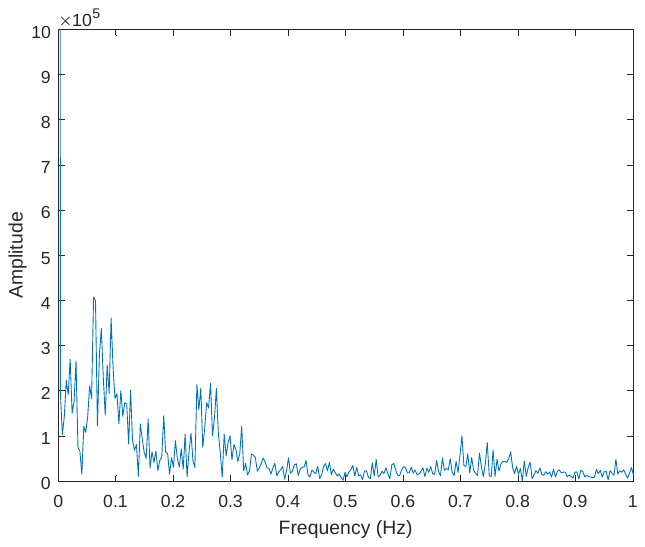 |
| 4 | 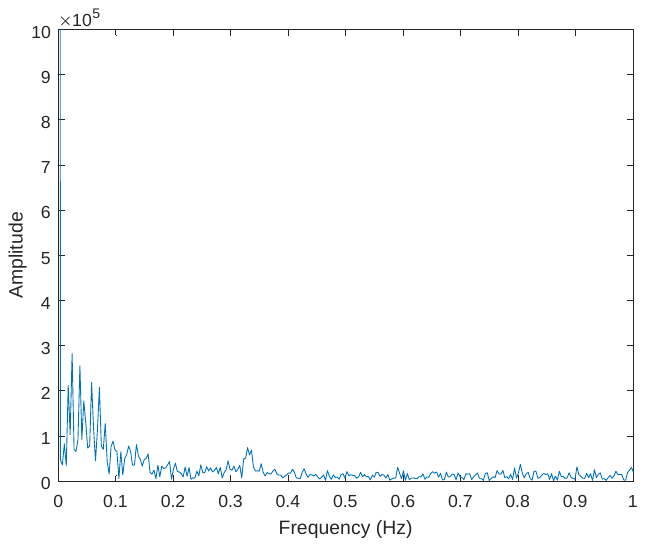 | 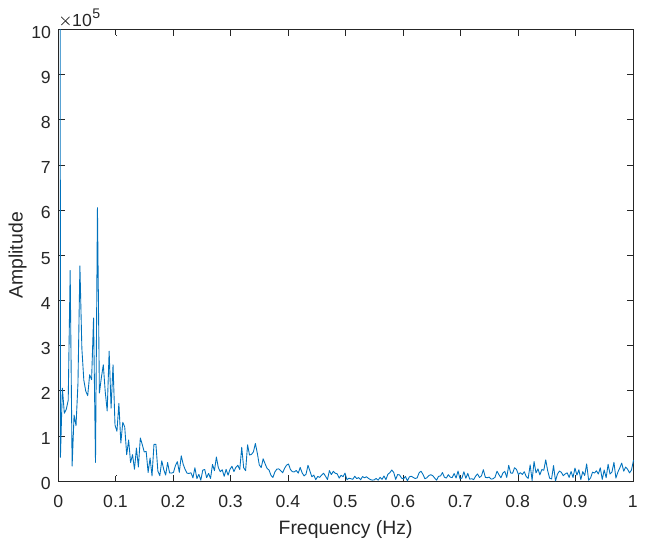 |
| 5 | 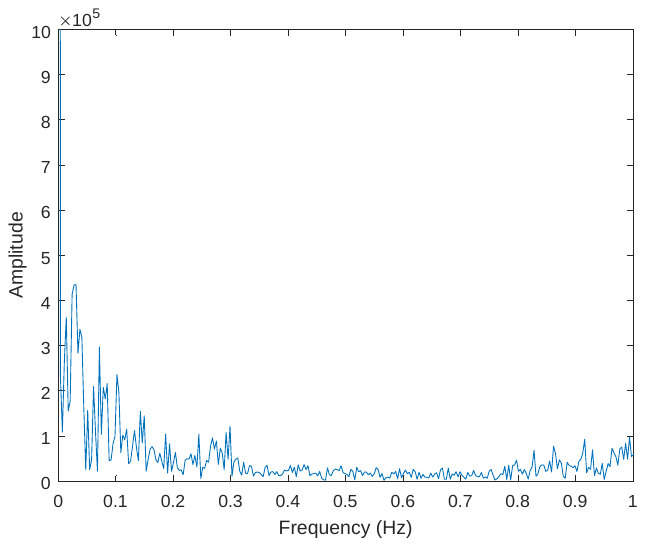 | 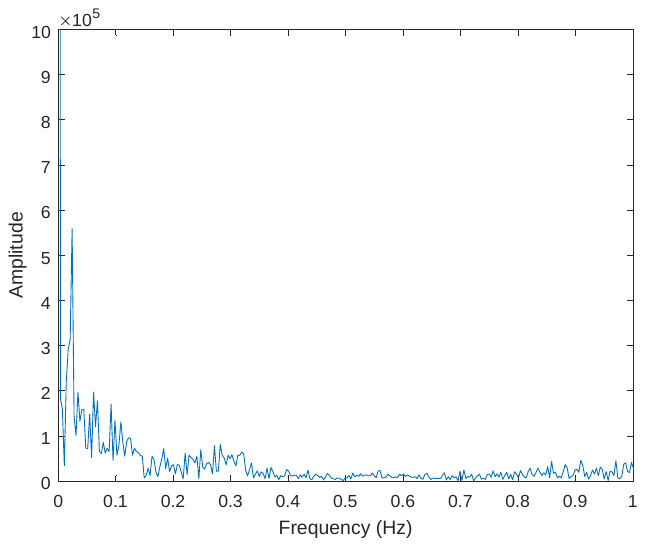 |
| 6 | 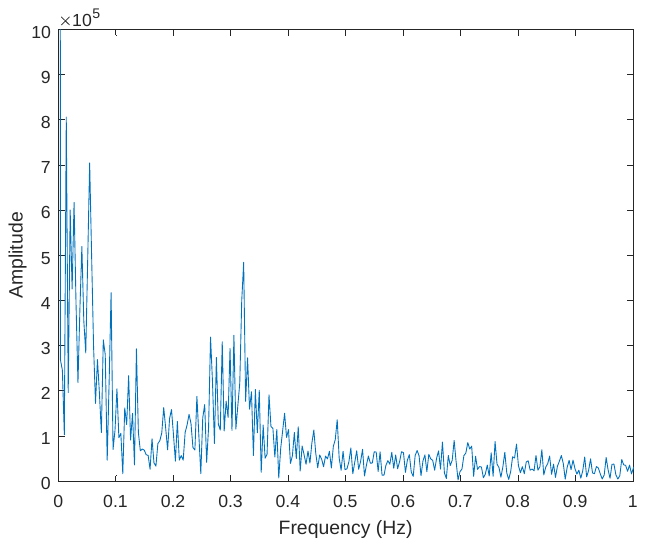 | 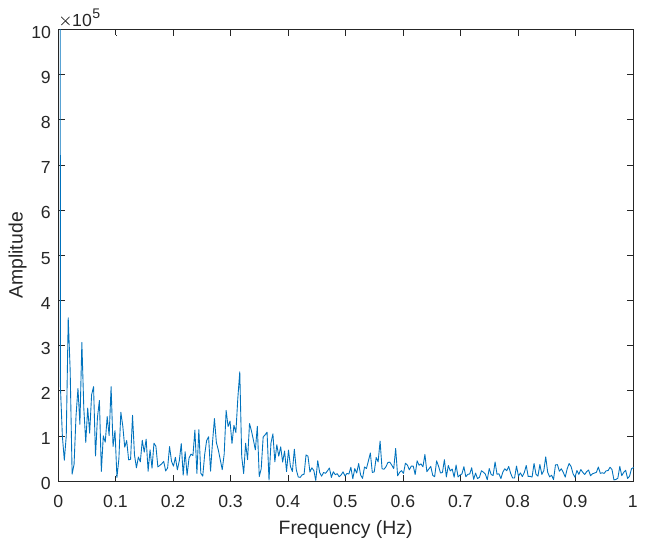 |
| 7 | 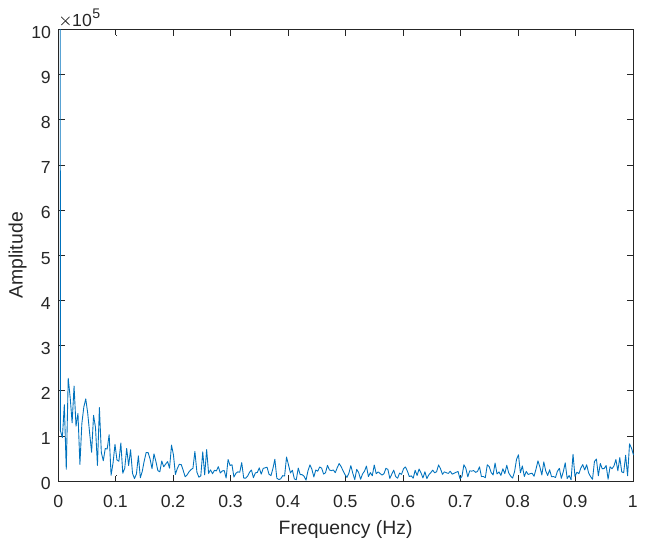 | 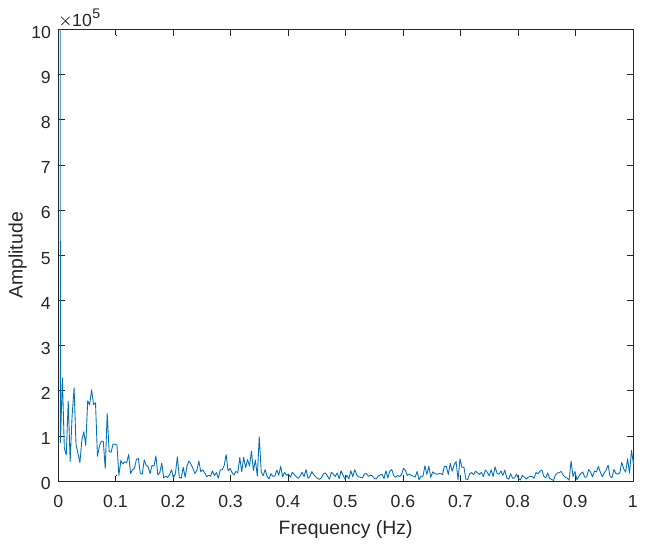 |
| 8 | 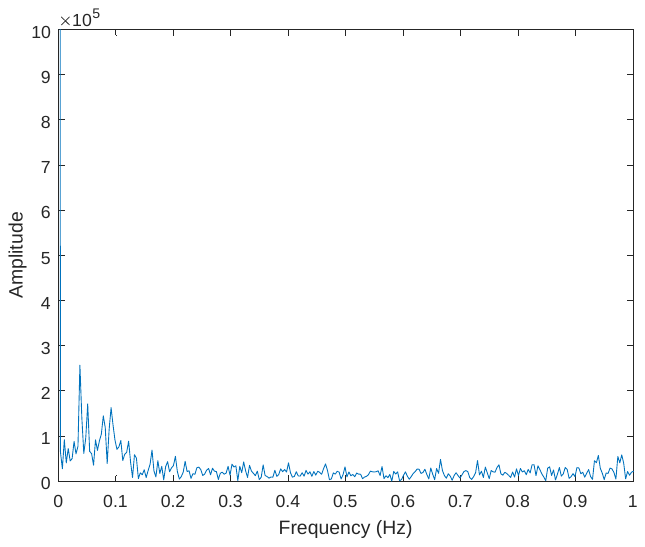 | 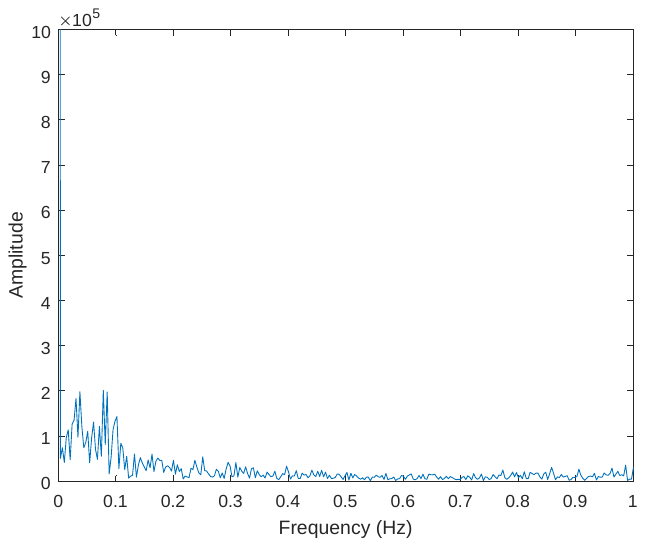 |
| 9 | 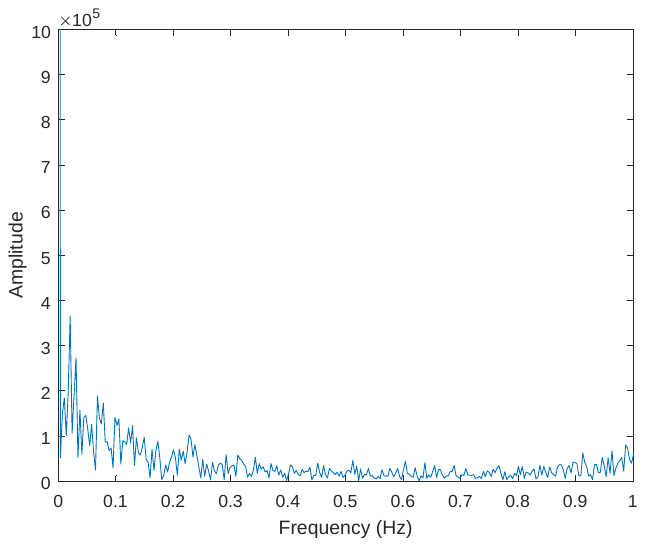 | 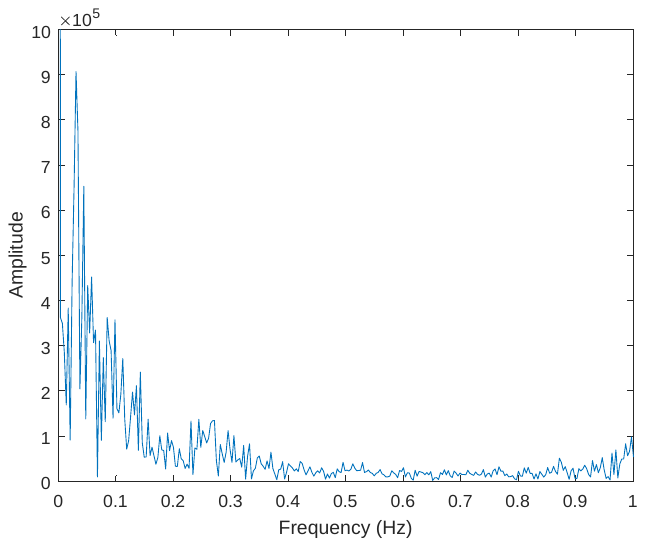 |
| 10 | 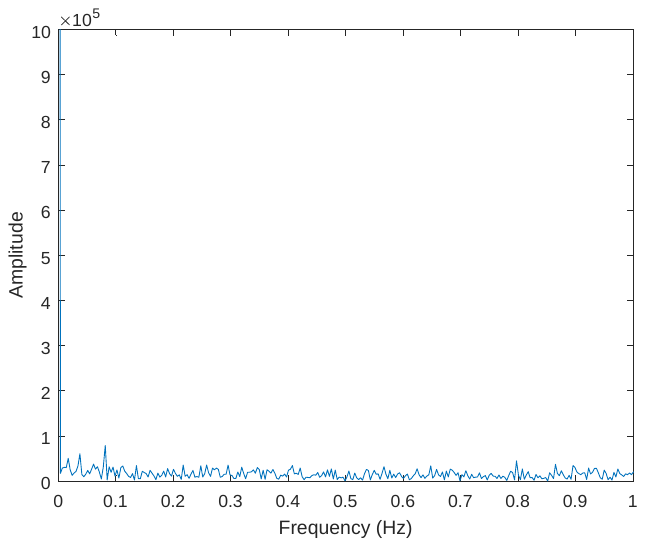 | 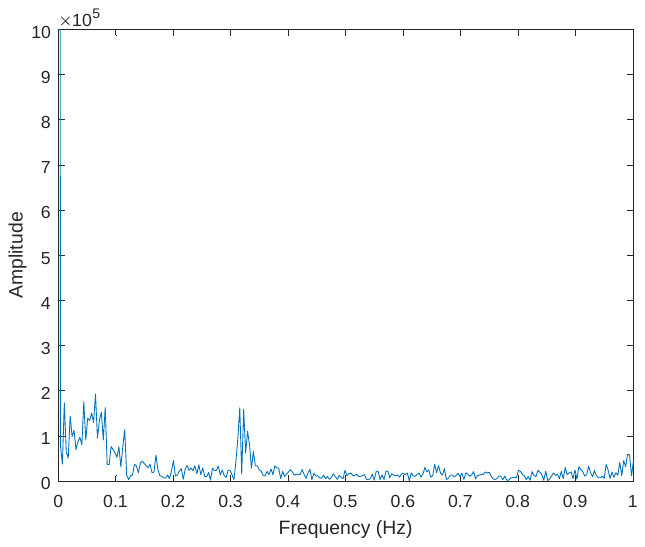 |
| Table S1. Power spectrums for CSF inflow fluctuation and outflow fluctuation signals. | | |

| participant | Spectrum for GMS brain | Spectrum for IJV neck |
| --- | --- | --- |
| 1 | 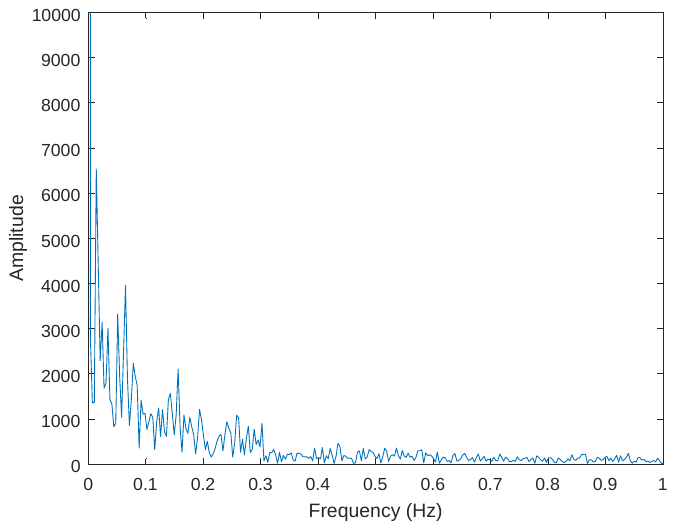 | 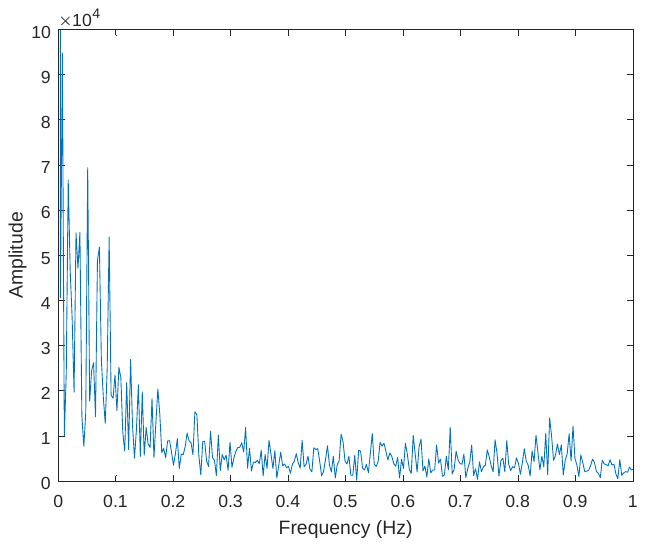 |
| 2 | 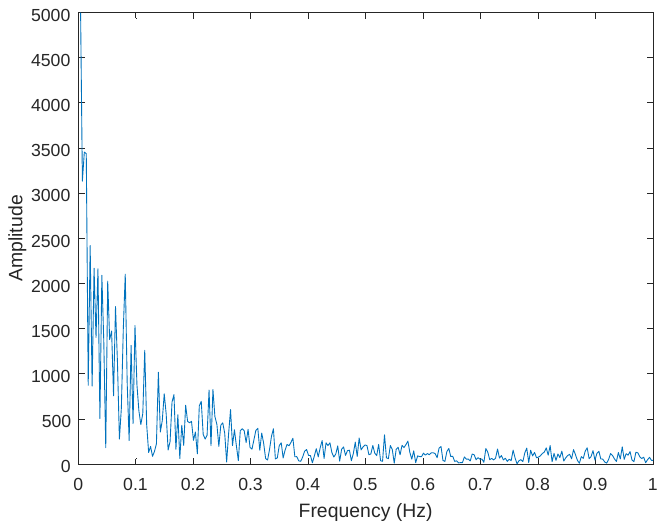 | 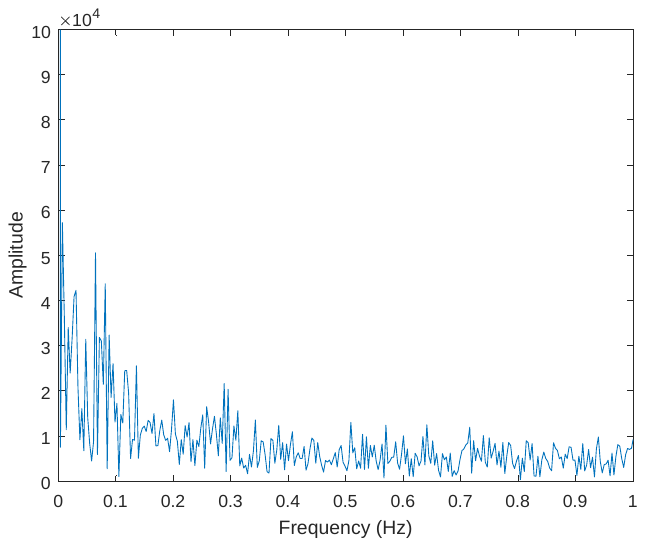 |
| 3 | 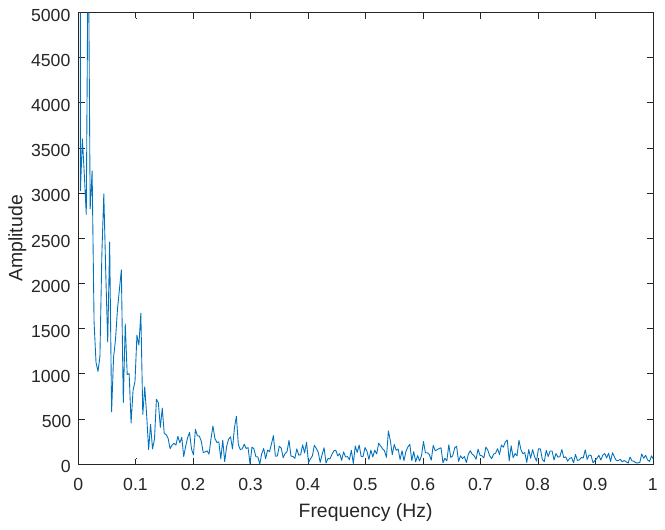 | 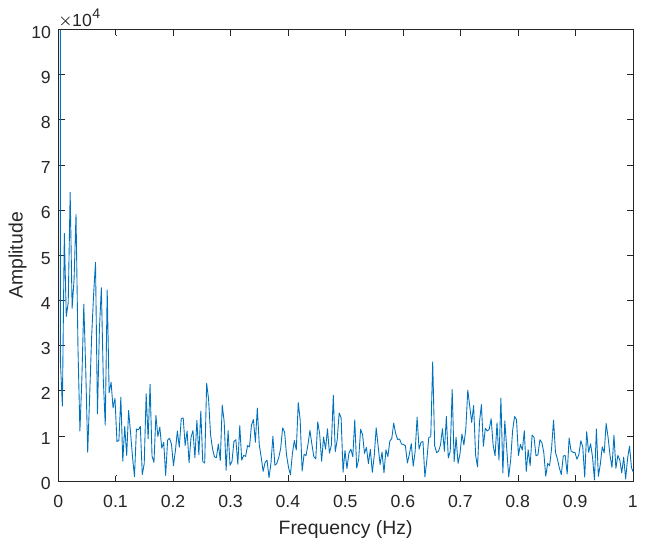 |
| 4 | 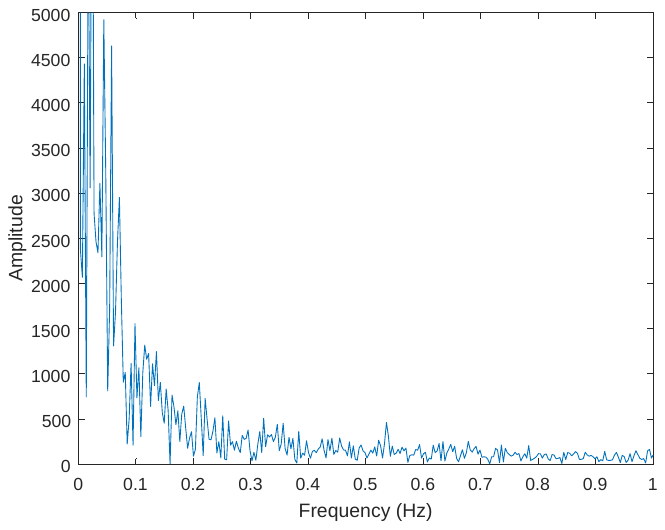 | 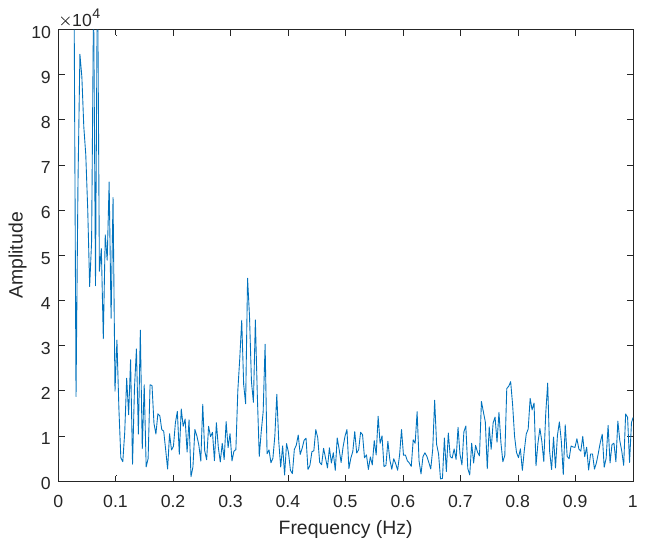 |
| 5 | 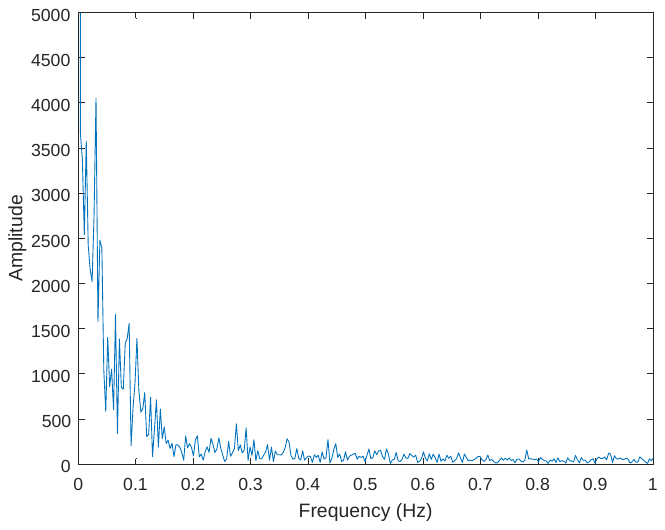 | 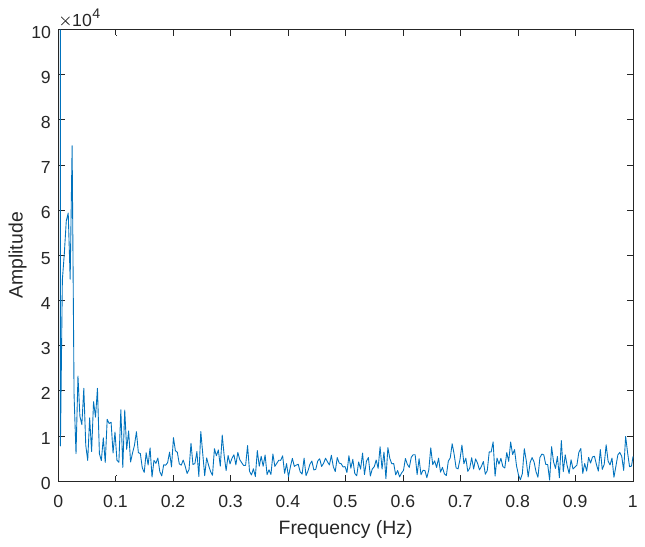 |
| 6 | 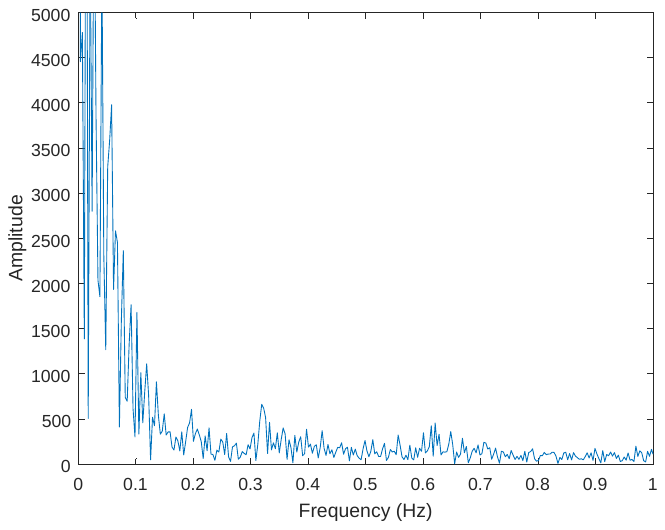 | 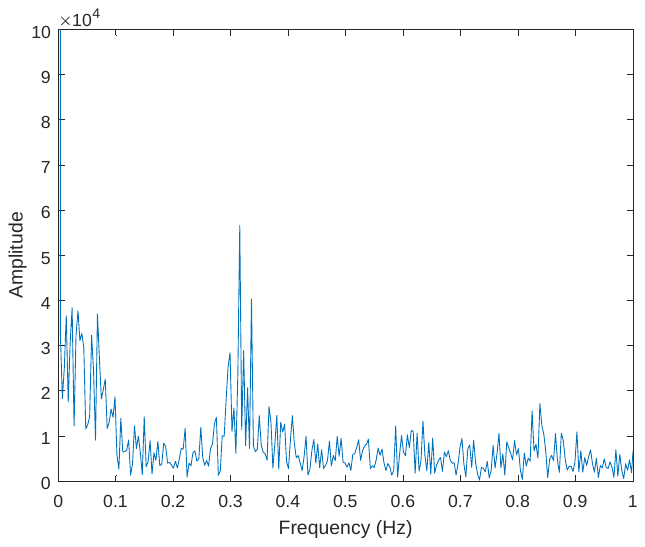 |
| 7 | 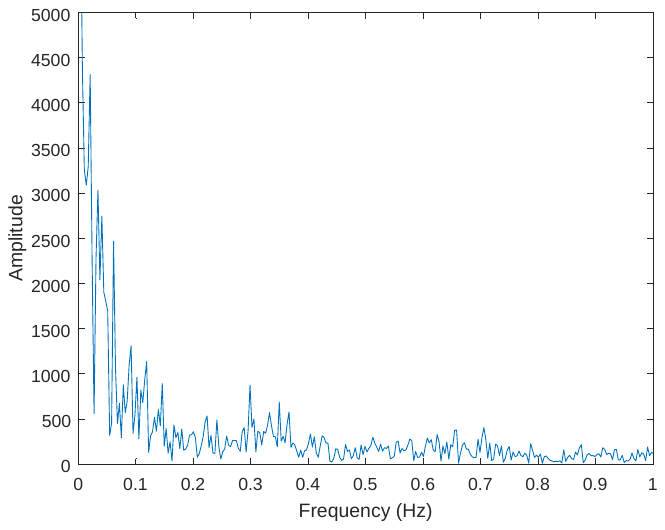 | 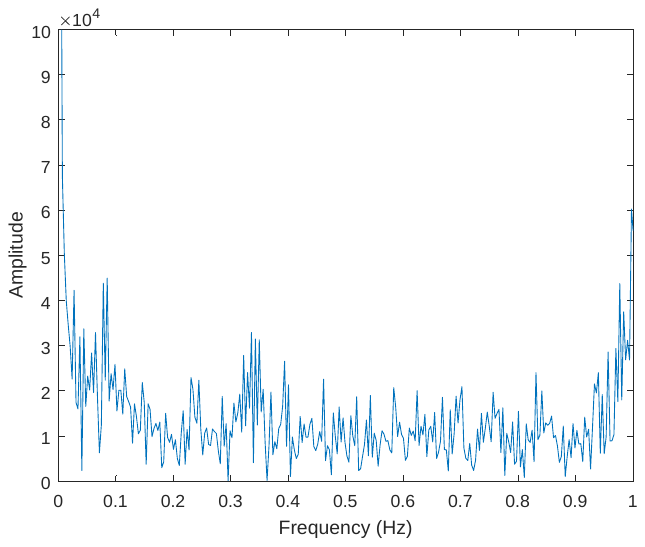 |
| 8 | 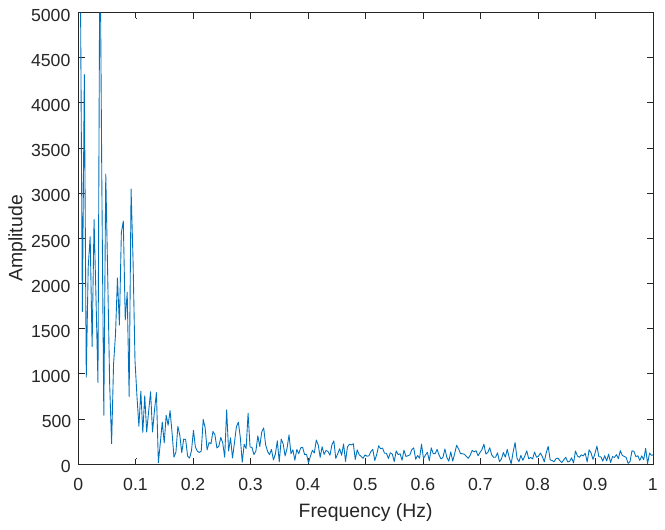 | 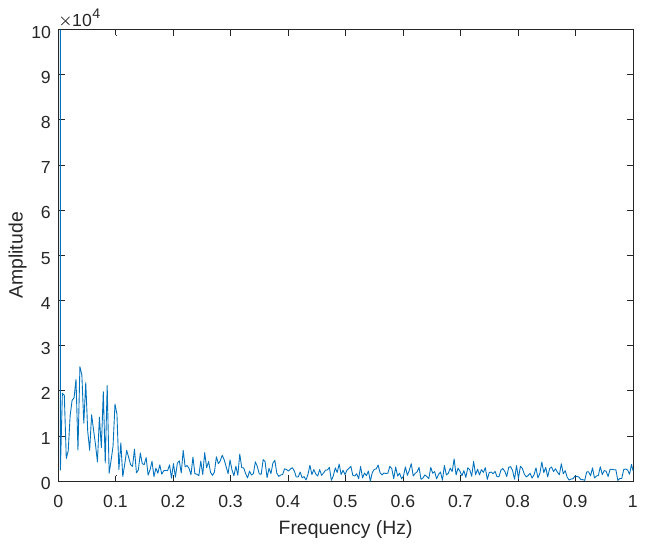 |
| 9 | 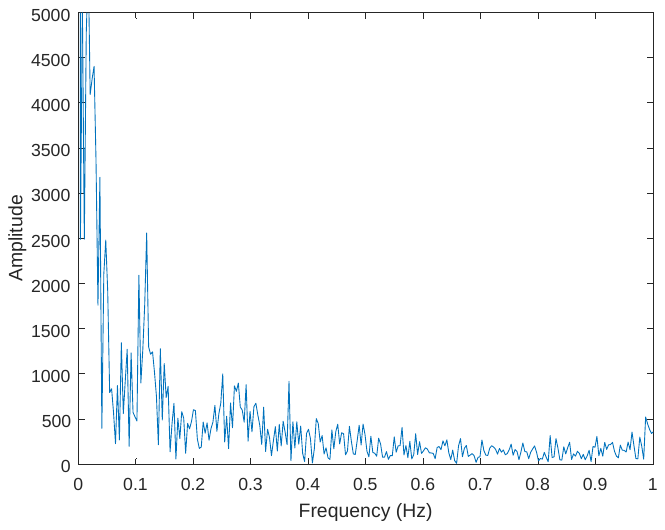 | 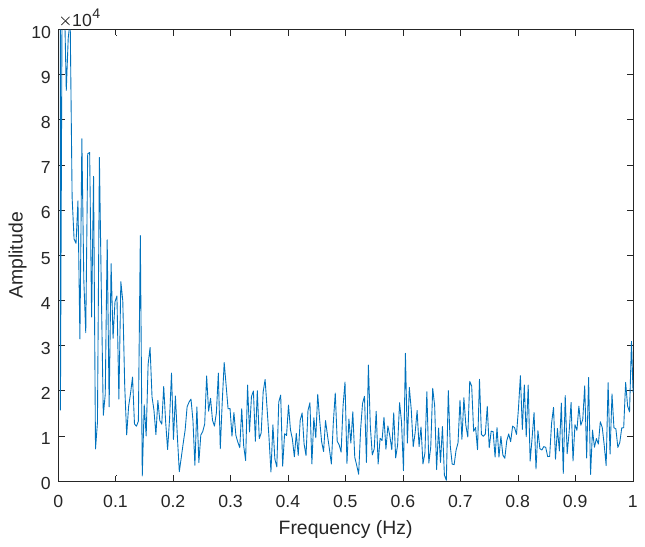 |
| 10 | 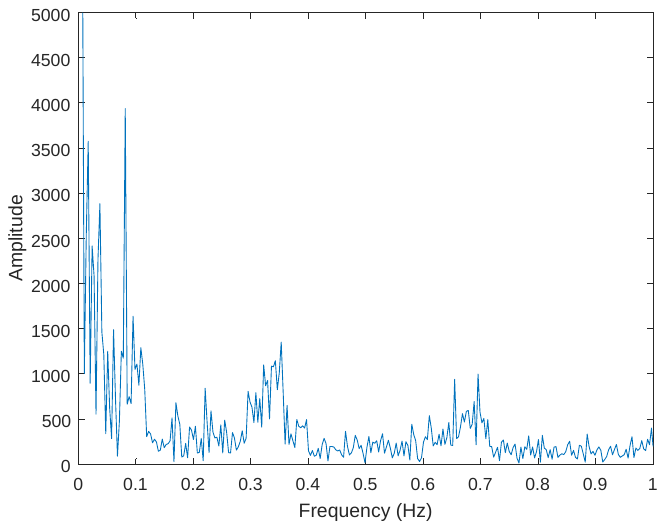 | 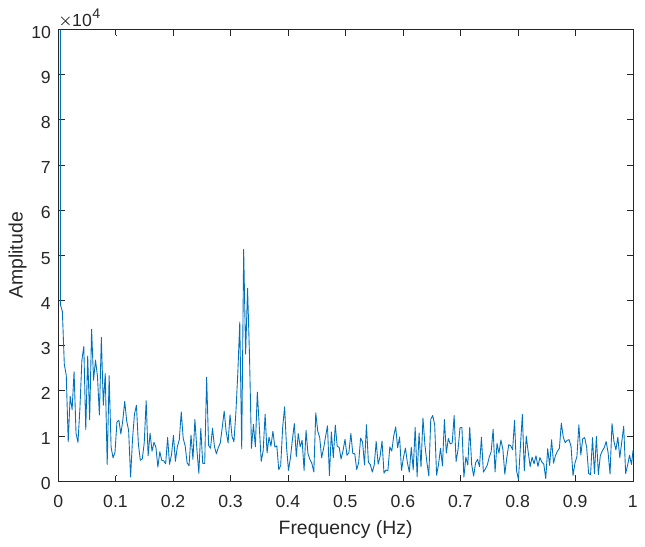 |
| Table S2. Power spectrums for GMS and IJV signal. | | |

| participant | positive-MCCC | lag (s) | negative-MCCC | lag (s) |
| --- | --- | --- | --- | --- |
| 1 | 0.5362 | -5.72 | -0.4973 | 3.08 |
| 2 | 0.5101 | -4.84 | -0.4023 | 3.52 |
| 3 | 0.6309 | -5.28 | -0.5487 | 3.52 |
| 4 | 0.6495 | -7.48 | -0.6544 | 2.64 |
| 5 | 0.5781 | -10.12 | -0.7128 | 3.08 |
| 6 | 0.6016 | -7.04 | -0.7103 | 3.96 |
| 7 | 0.6309 | -8.36 | -0.6232 | 3.08 |
| 8 | 0.645 | -5.28 | -0.6794 | 3.08 |
| 9 | 0.6426 | -9.68 | -0.5335 | 4.4 |
| 10 | 0.5997 | -5.28 | -0.4288 | 2.2 |
| Table S3. Separately calculated positive MCCCs and negative MCCCs and corresponding time delays between $\mathrm{GMS}$ and CSF inflow fluctuation. | | | | |

| participant | positive-MCCC | lag (s) | negative-MCCC | lag (s) |
| --- | --- | --- | --- | --- |
| 1 | 0.6362 | -3.52 | -0.6091 | 3.08 |
| 2 | 0.5996 | -5.28 | -0.5989 | 4.84 |
| 3 | 0.7553 | -3.96 | -0.7531 | 3.52 |
| 4 | 0.6718 | -4.84 | -0.669 | 4.4 |
| 5 | 0.6772 | -4.84 | -0.6874 | 4.4 |
| 6 | 0.6455 | -4.4 | -0.6045 | 4.4 |
| 7 | 0.7316 | -4.84 | -0.7335 | 4.4 |
| 8 | 0.6456 | -4.4 | -0.6368 | 3.96 |
| 9 | 0.6517 | -3.96 | -0.6668 | 3.52 |
| 10 | 0.6294 | -3.96 | -0.6725 | 3.52 |
| Table S4. Separately calculated positive MCCCs and negative MCCCs and corresponding time delays $\mathrm{GMS}$ and the $\frac{d}{\mathrm{dt}}(GMS)$. | | | | |

**CSF flux assessment**

In this study and previous research, CSF flowed in both directions through the fourth ventricle ^5-18^. To roughly assess the CSF flux, we calculated the areas under 1) the CSF inflow fluctuation (from brain scan, Figure S1.a red) and 2) CSF outflow fluctuation (from neck scan, Figure S1.a blue) for each participant (two scans are back-to-back with participant being in the same position, Figure S1.b). The difference (Figure S1.c) represents the net amount of CSF flux during the time of scan for each participant. It is known that the CSF inflow and outflow fluctuation were measured consecutively in two separated scans. Thus, the calculation of the CSF flux is not strictly accurate and should be treated as an estimate. We have no reason to believe that CSF flow fluctuation would change drastically in the short time between scans; therefore, it was reasonable to assume they could be combined to generate an estimate.

No consistent CSF flux direction was detected among participants, indicating the CSF flux was not substantial during the short scan time (i.e., minutes). It is known that adults only secret 25 ml CSF/hour in the brain (mostly from the choroid plexuses of the lateral ventricles) ^19^. The secretion creates a pressure that dictates the direction of the CSF flow fluctuation through the ventricular system to the subarachnoid space (CSF flux in one direction). However, due to the small amount of CSF, the CSF flux would be hard to detect during the short period of time with our imaging method.

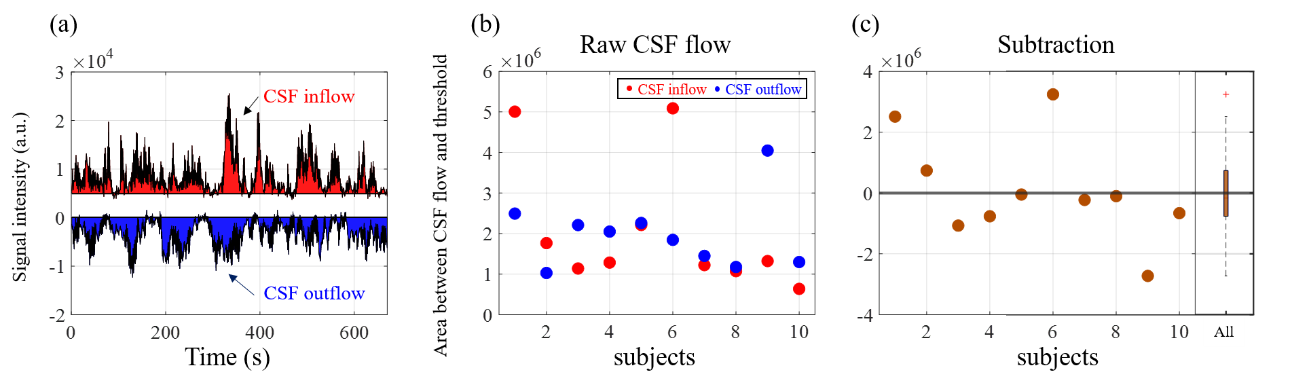

Figure S1. CSF flux assessment (a) Area above/below CSF fluctuations in in/out direction was calculated as flux within one scan. (b) Summarize of in/out flux for each participant. (3) CSF flux was assessed by subtracting influx from outflux.

**High correlations were found between brain (GMS) and big blood vessels in the neck (IJV)**

Two participants were scanned, and the similar protocol was applied while covering from the brain to neck (TR =1.13s, 90 slices, 360 volumes to make the full coverage). GMS signals of fMRI was calculated by averaging fMRI signal in the brain (Figure S2, orange), while IJV signal was calculated by averaging fMRI signal in the identified IJV mask (Figure S2, light-blue). Both GMS and IJV signals were detrended and band-pass filtered (0.01-0.1Hz) according to previous publication^20^. In Figure S3, we validated that the low frequency oscillation in the IJV is correlated with GMS (participant 1: 0.46; participant 2: 0.69) with few second delay (participant 1: 3.39s; participant 2: 4.52s).

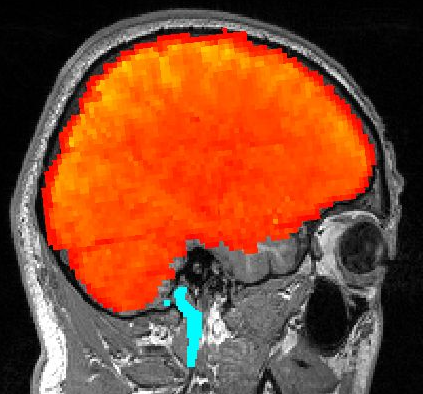

Figure S2. Region of interests of GMS and IJV.

| Participant1 | Participant2 |
| --- | --- |
| 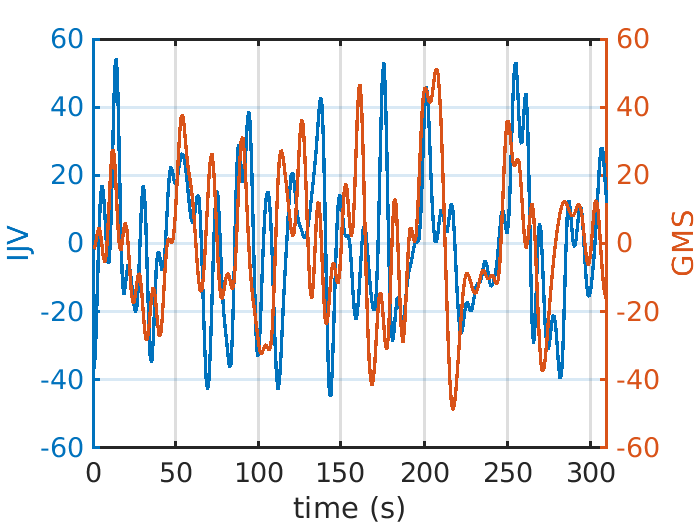 | 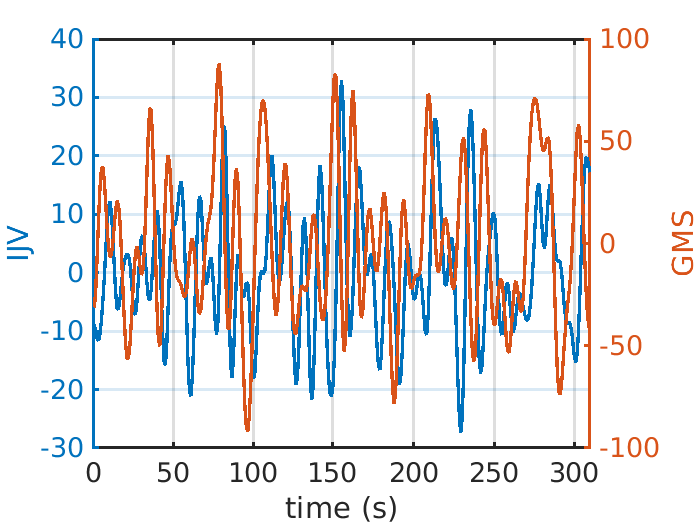 |
| Figure S3. Two participants’ GMS and IJV signals. | |

**Adaption of our original model with pulsatile effect on CSF fluctuations**

We also adapted our model in Figure 1 (see in manuscript) by adding the pulsatile cardiac oscillations. Thus, the model included both pulsatile and slow arteriolar oscillations on the blood vessels (Figure S4), in which, the faster and smaller pulsatile changes were added to the slow, bigger low frequency changes. From the model, we can see that 1) inflow fluctuation will be continuous during the slow transition period. 2) The pulsations during this period will modulate the inflow fluctuation speed (not reverse flow direction), thus full scale of the derivative pulsatile signal is “recorded” by the fMRI. However, for the pulsations happening outside this period, they induced CSF flow fluctuation in both directions. Therefore, only partial signal (positive derivative pulsatile signal) was “recorded”.

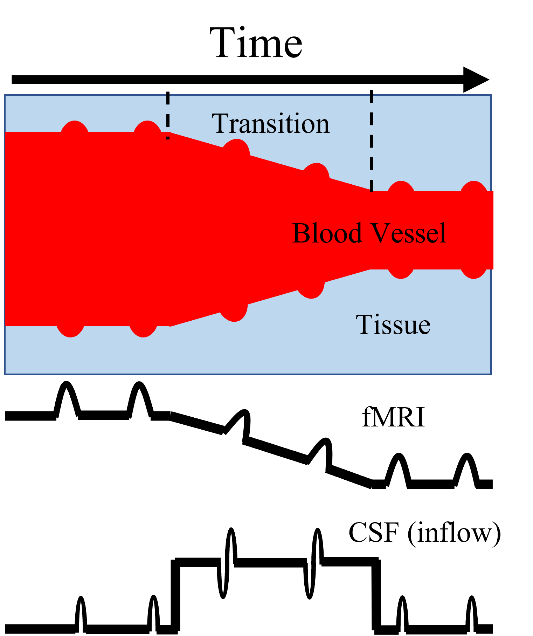

Figure S4. Adaptive model with pulsatile effect on CSF fluctuations.

**Heartbeat and limited cardiac effect on low frequency oscillations**

Power spectra for each participant’s CSF in/out flow fluctuation was provided in Table S1. However, heart rate can only be assessed from one participant (0.87 Hz), heart rates for other participants were above the Nyquist frequency of the acquisition (1.13 Hz). We conducted a quick assessment on the CSF data that fully sampled the cardiac pulsation. We tried to estimate the impact of heartbeat on the low frequency (0.01-0.1Hz) via aliasing. A hundred high frequency (1-2Hz) signals (mimicking heartbeat signal) with 4Hz sampling rate and one third amplitude of low frequency signal (the ratio is similar to what we found in one of the participants’ amplitude ratio between heartrate and low frequency component) were simulated to test their influence on a simulated low frequency signal. First, high frequency signal was combined with low frequency signal and resampled in 2.27Hz (same as our fMRI study). Second, the combined signals will be filtered (0.01-0.1Hz). Then, the filtered signal (with aliased heartbeat signal) was then correlated with original low frequency signal. In Figure S5, these correlations remain high (>0.98), which indicated high frequency (heartrate) signals could only explain limited variance in low frequency signal through aliasing. The result is consistent with our previous publication, in which, we showed that aliased cardiac pulsations signals were not the main component of LFOs signals in fMRI ^21^.

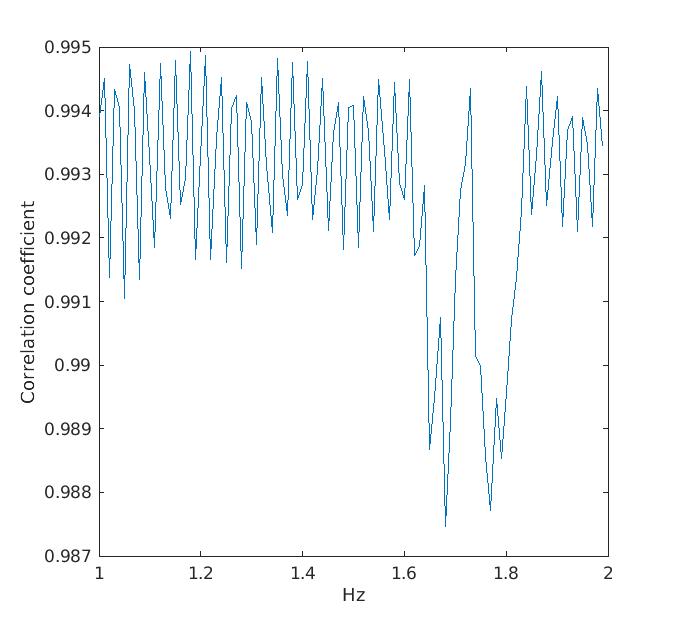

Figure S5. Simulation results for cardiac effect on low frequency signal. X-axis: one hundred high frequency signals (1-2Hz); Y-axis: correlation of original low frequency signal and low frequency signal mixed with high frequency signals.

**Investigation of motion effect on CSF fluctuations**

To find out if our observations are due to motion effects, we calculated the correlation between six motion parameters and CSF flow fluctuation-related signals. Our results show the correlations were low (mean value lower than 0.2) between motion parameters and CSF signal for all the participants under filtered (band-pass filtering between 0.01-0.1Hz) condition (Table S5).

|  | brain | | | | | | neck | | | | | |
| --- | --- | --- | --- | --- | --- | --- | --- | --- | --- | --- | --- | --- |
| participant | $r_{x}$ | $r_{y}$ | $r_{z}$ | $t_{x}$ | $t_{y}$ | $t_{z}$ | $r_{x}$ | $r_{y}$ | $r_{z}$ | $t_{x}$ | $t_{y}$ | $t_{z}$ |
| 1 | 0.14 | 0.04 | 0.03 | 0.00 | -0.04 | -0.21 | -0.14 | -0.09 | -0.11 | -0.10 | 0.00 | 0.12 |
| 2 | 0.25 | 0.18 | -0.07 | -0.15 | 0.03 | -0.20 | 0.12 | 0.19 | -0.21 | -0.16 | -0.25 | 0.12 |
| 3 | -0.14 | -0.06 | -0.02 | -0.01 | -0.08 | 0.01 | -0.02 | -0.01 | 0.05 | 0.05 | -0.14 | -0.04 |
| 4 | 0.25 | -0.06 | -0.09 | -0.13 | -0.17 | 0.05 | 0.02 | -0.03 | 0.15 | 0.13 | -0.06 | -0.07 |
| 5 | -0.01 | 0.05 | 0.12 | -0.07 | -0.14 | -0.07 | 0.11 | 0.05 | 0.14 | 0.05 | -0.13 | -0.08 |
| 6 | 0.25 | -0.12 | -0.06 | 0.10 | -0.14 | -0.22 | 0.29 | -0.26 | 0.06 | 0.14 | -0.26 | -0.20 |
| 7 | -0.01 | -0.04 | -0.23 | -0.05 | -0.03 | 0.23 | 0.26 | -0.13 | 0.15 | 0.12 | -0.16 | -0.18 |
| 8 | 0.08 | 0.08 | -0.01 | -0.10 | -0.10 | 0.09 | -0.10 | 0.22 | 0.09 | 0.01 | 0.01 | 0.25 |
| 9 | 0.10 | 0.06 | 0.06 | -0.20 | -0.10 | -0.06 | -0.04 | 0.01 | 0.09 | 0.09 | 0.00 | -0.12 |
| 10 | 0.05 | -0.02 | 0.01 | -0.02 | -0.13 | -0.06 | 0.35 | 0.00 | -0.19 | 0.05 | -0.29 | -0.18 |
| mean | 0.10 | 0.01 | -0.03 | -0.06 | -0.09 | -0.04 | 0.08 | -0.01 | 0.02 | 0.04 | -0.13 | -0.04 |
| std | 0.13 | 0.09 | 0.10 | 0.09 | 0.06 | 0.15 | 0.17 | 0.14 | 0.14 | 0.10 | 0.11 | 0.15 |
| Table S5: Correlations between mean motion parameters and CSF fluctuations after band-pass filtering (0.01-0.1Hz). $r_{x-z}$: MCFLIRT estimated rotations. $t_{x-z}$: MCFLIRT estimated translations. | | | | | | | | | | | | |

Also, we accessed the signal which is one voxel above/below the original selected CSF voxel, which is motion corrected, and called it CSF signal in the second slice. Six out of ten participants showed that this CSF signals in the second slices are still correlated (r >0.3) with original CSF signals in the first slice. Most importantly, CSF signals in the second slice still lag the $\frac{d}{\mathrm{dt}}(GMS)$. For the other 4 participants, CSF signal in the second slice did not have high correlation (r<0.3) with original CSF signal. Also, their time delays to the $\frac{d}{\mathrm{dt}}(GMS)$ are variable.

To further investigate motion effect on CSF signal, CSF signal in the second slice (one voxel above (brain) or below (neck) of original CSF signal) was examined. We observed lower sensitivity from the CSF signal in the second slice. In addition to the obvious reason that inflow effect is weakened as particles move further into the scan volume (it depends on the flow speed too), partial volume effects might also contribute to the observation. The areas of the fourth ventricle are generally wider in the first slice than the second slice. Therefore, the chance of having non-CSF voxels is higher in the second slice. The results were presented in Table S6 and S7. In summary, we did not find the motion artefact affected the CSF signal from the first slice.

| Brain | | | |
| --- | --- | --- | --- |
| participant | corr_CSF_CSFv2 | xcorr | lag (s) |
| 1 | 0.75 | -0.78 | -2.2 |
| 2 | 0.06 | 0.25 | -13.2 |
| 3 | 0.84 | -0.75 | -1.32 |
| 4 | 0.07 | -0.24 | 6.16 |
| 5 | 0.49 | -0.45 | -4.4 |
| 6 | 0.86 | -0.85 | -1.32 |
| 7 | 0.59 | -0.45 | -6.16 |
| 8 | 0.05 | -0.22 | 14.96 |
| 9 | 0.68 | -0.44 | -5.72 |
| 10 | 0.23 | -0.25 | -1.32 |
| mean | 0.46 | -0.42 | -1.45 |
| std | 0.31 | 0.31 | 7.18 |
| Table S6. CSF fluctuations in second slice for brain scan. First row: correlation between CSF fluctuations in first slice (original ROI) and second slice (slice with motion correction). Second and third rows: cross-correlation results between CSF fluctuations in second slice and $\frac{d}{\mathrm{dt}}(GMS)$. | | | |

| Neck | | | |
| --- | --- | --- | --- |
| participant | corr_CSF_CSFv2 | xcorr | lag (s) |
| 1 | 0.96 | 0.62 | 3.52 |
| 2 | 0.08 | -0.37 | -3.08 |
| 3 | 0.92 | 0.76 | 1.76 |
| 4 | 0.38 | 0.30 | -1.76 |
| 5 | 0.43 | 0.34 | 1.76 |
| 6 | 0.78 | 0.20 | 10.56 |
| 7 | 0.68 | 0.31 | 0 |
| 8 | 0.01 | -0.34 | -7.04 |
| 9 | 0.92 | 0.58 | 2.2 |
| 10 | 0.18 | -0.18 | 12.76 |
| mean | 0.53 | 0.22 | 2.07 |
| std | 0.35 | 0.38 | 5.63 |
| Table S7. CSF fluctuations in second slice for neck scan. First row: correlation between CSF fluctuations in first slice (original ROI) and second slice (slice with motion correction). Second and third rows: cross-correlation results between CSF fluctuations in second slice and $\frac{d}{\mathrm{dt}}(IJV)$. | | | |

Together, these results demonstrate that the main results in the paper are not due to motion effects.
